# The impact of temperature-induced vertebral anomalies on C-start swimming performance in Astyanax mexicanus (Teleostei: Characidae)

**DOI:** 10.64898/2026.08.06.743195

**Authors:** Kaleigh M. Arnold, Winer Daniel Reyes-Corral, Owen Howard, Christian Graca, Windsor E. Aguirre

## Abstract

This study investigates the impact temperature-induced vertebral anomalies have on the C-start escape response of *Astyanax mexicanus*, a model species in evolutionary developmental biology. Employing three temperature treatments to induce varying degrees of skeletal anomalies, we assessed their effects on key swimming performance metrics including, C-start time, curvature coefficient, head displacement distance, and displacement velocity. Through the use of linear mixed models and generalized linear mixed models, our results reveal that specific anomalies such as vertebral fusions and anomalous haemal and neural spines affected the curving ability of C-start escape responses. However, these did not negatively impact other performance parameters, with velocity, distance, and response time showing no significant impacts from any anomaly types, when assessed individually. This suggests a complex interplay between structural deformities and compensatory physiological mechanisms that maintain functional performance. Other variables measured had a stronger and significant impact on swimming performance, including standard length, vertebral number, and temperature treatment, which influenced escape speed, curving ability, and overall locomotor performance. Our findings challenge conventional perceptions about the debilitating impact of vertebral anomalies, indicating that many affected fish can still effectively perform escape maneuvers critical for survival.

## Introduction

Skeletal anomalies in fish can result from factors such as poor water quality, nutritional imbalances, exposure to heavy metal pollutants, changes in water conditions, temperature fluctuations, and underlying epigenetic factors (1–5). Research on skeletal anomalies has primarily focused on economically valuable temperate fish species cultivated in Europe and North America due to issues anomalies present during filet production. (6–8). As wild fish populations in the Neotropics and other tropical regions face unique climatic changes and increasing environmental pollutants, it is essential to expand research to include Neotropical freshwater fish (NFF) species (2,6).

Among skeletal anomalies, those affecting the vertebral column, such as curvatures, fusions, and compressions, are commonly observed and can significantly impact fish fitness, particularly in terms of locomotion (7,9). Anomalies in the vertebral column can impair a fish’s ability to effectively escape from predators and catch food, leading to strong selective pressures against anomalous fish in the wild (2,10). Reports of sexually mature fish with vertebral or skeletal anomalies in the wild are relatively limited (although increasing), likely due to the high mortality rates of deformed embryos and compromised physiological capabilities of older deformed fish (3,10–12). The ability to escape predators may be compromised in fish with vertebral anomalies.

The C-start is a startle response in which a fish rapidly bends its body into a characteristic “C” shape following a sudden stimulus and plays a key role in predator avoidance in fishes. It consists of two primary stages: a preparatory stage, where the fish quickly curls its body into a “C” shape by bringing its head and tail together while turning its body away from the initial stimulus, and a propulsion stage, where the fish escapes using the force generated as it straightens its body to flee from a threat (13–15) . Because vertebrae play a central role in the rapid bending of the body during the C-start response, this behavior may be particularly vulnerable to the effects of vertebral anomalies. However, there is currently limited research on how such anomalies affect escape response like the C-start.

Here, we examine how temperature-induced vertebral anomalies impact the C-start escape response in laboratory-reared surface morphotypes of *Astyanax mexicanus*. To achieve this, we established three different treatments, varying in rearing temperature and timing, specifically designed to induce these anomalies (5,16,17). We will assess the influence of the type, amount, and placement of vertebral anomalies on various parameters of C-start swimming performance, including the C-start time, the curvature coefficient (CC), head displacement distance (HDD), and displacement velocity (DV). *Astyanax mexicanus* has emerged as a model organism in evolutionary developmental biology in the last decade and is a member of the hyper-diverse order Characiformes, which is the second largest taxonomic order in the Neotropics (18–23). We predict that the type, amount, and placement of vertebral anomalies will negatively impact swimming parameters, resulting in decreased escape performance. By examining how temperature influences the development of vertebral anomalies and how these anomalies affect escape performance, we aim to broaden our understanding of the functional consequences of skeletal anomalies and their potential implications for fitness.

## Methods

### Breeding and fish care

All experimental procedures involving animals were conducted in accordance with Protocol No. IACUC-2022-640 approved by the DePaul University Institutional Animal Care and Use Committee. Fish used in this study originated from an established breeding colony of surface morph *Astyanax mexicanus* at DePaul University. The breeding colony was derived in 2014 from 4th generation descendants of wild-caught *A. mexicanus* individuals from the Rio Grande River, Texas National Park, maintained by the Jefferey Lab at the University of Maryland. Male breeders were kept in 5.5-gallon community tanks, while female breeders were separated and kept in groups in 20-gallon tanks.

A total of 10 crosses were performed in July and August of 2022 to obtain approximately 300 fish for the experiment (10 fish per each of the three temperature treatments resulting in 30 fish for each of the 10 crosses). For the crosses, ten new 5.5-gallon tanks were set up with sponge filtered aerators and aquarium heaters to maintain a stable temperature of 25 °C, promoting breeding. A single pair of male and female breeders were placed in each 5.5-gallon tank early in the morning (by 8 am) to acclimate and spend the day together. The breeding pairs consisted of eight pairs of 1-year-old fish and two pairs of 5-year-old fish. Two weeks prior to the initial crosses, females were fed a diet of frozen adult brine shrimp twice daily to prepare for breeding. The lights were turned off at 6pm on the same day that the crosses were set up, and the tanks were monitored approximately every 30 minutes to one hour for eggs.

For the crosses that spawned, fertilized eggs were collected between 10 pm and 2 am and distributed evenly across three temperature treatments: 21±1°C (constant or C), 30±1°C for one month transitioning to 21±1°C (short-term or ST), and 30±1°C for three months transitioning to 21±1°C (long-term or LT). Eggs in the low temperature treatment were housed in petri dishes exposed to the ambient facility temperature of 21±1°C, while eggs in the elevated temperature treatments were reared in petri dishes in incubators set to 30±1°C. A diluted 1:20 methylene blue solution was added to each petri dish to prevent fungal growth. The solution was replenished after every water change until fish hatched.

After two weeks in the petri dishes, fish were transferred to 5.5-gallon tanks. Aquarium heaters were used in the tanks with the elevated 30±1°C temperature treatments. The tanks were filled with conditioned water following DePaul’s animal care facility guidelines, including sponge filtered aerators connected to external air pumps. Fish were fed live brine shrimp twice daily in petri dishes and transitioned to a diet of powdered Tetramine flakes and decapsulated brine shrimp eggs for two months before being fed only Tetramine flakes. Rotifers were periodically added to the tanks for enrichment. Weekly water changes of 25-33% were performed, and water quality parameters (ammonia, nitrate, alkalinity, pH, and hardness) were regularly monitored. Temperature was continuously monitored to ensure stable environmental conditions. To transition the fish from the elevated temperature to the low temperature, a 48-hour period at 25±1°C was implemented to prevent shock. After the transitional period, the heaters were removed, and the water temperature was equilibrated at 21±1°C. Fish were allowed to grow until the time of swimming performance trials, which began when the fish were approximately four months old. All fish were tested at a temperature of 21±1°C, after being acclimated for at least one week.

### Swimming Performance Trials

Swimming performance trials were conducted from December 2022 to January 2023 in the DePaul animal care facility. Fish were tested within a six-week timeframe. High-speed swimming performance videos were recorded using a Basler 504k high-speed area scan camera with a Nikon AF Micro Nikkor 60mm f/2.8D lens and the EPIX © XCAP™ Image Processing Software system. Videos were recorded at 1000 frames per second (fps) with an aperture of 2.8.

The testing setup included a mirror positioned at a 45° angle toward the camera, displaying the bottom half of the testing vessel. A clear acrylic bowl with a height of 12cm and a diameter of 24.3cm served as the testing vessel where reactions were recorded. The water level was maintained at 3cm to prevent vertical swimming movements that could not be recorded. Two light sources were positioned toward the testing vessel, one from above and one from below. A reservoir of fresh aerated water was kept near the testing vessel at a stable temperature of 21±1 °C, allowing for consistent water changes in 50mL increments to maintain oxygen and temperature levels during testing.

C-start swimming performance was recorded for each fish, consisting of two trials. Fish were allowed to acclimate in the testing vessel for 5 minutes before each trial, and the duration in the vessel was limited to a maximum of 20 minutes. To elicit escape responses, a 1.25g/2cm pebble was used as a stimulus and dropped to the left side of the fish. Recordings started just before the pebble hit the water and ended after the first full tail beat. A trial was considered successful if the fish exhibited both stages of the C-start response. If a fish was unreactive during a trial, a secondary trial was conducted after a five-minute interval, with a maximum of four attempted trials to avoid stress or acclimation to the stimulus. After two trials were recorded, fish were euthanized in an ice water bath and then stored in 10% formalin for at least 48 hours before being transferred to 70% ethanol for long-term storage. Standard length (SL) was measured from the formalin fixed specimens starting at the tip of the fish’s head to the posterior end of the caudal peduncle.

### Swimming Performance Parameters

Several swimming performance parameters were measured using Tracker: Video Analysis and Model Tool (Brown, 2024, Fig 1). C-start time was measured in seconds (C-sT) starting from the beginning of stage 1, when the head started to move away from the stimulus to the end of the tailbeat in stage 2. The Curvature Coefficient (CC) was calculated by dividing the maximum curvature (MC) by the SL and then subtracted from 1, (1-(MC/SL)), where MC is defined as the shortest distance between the head and the tail when the body was bent during the C-start response in stage . Larger values for the CC indicate a greater curvature of the body per unit size. Initial and final head coordinates were recorded to calculate head displacement distance (HDD) in cm using the distance formula (d=√((x2 – x1)² + (y2 – y1)²)). Displacement velocity (DV) was measured as the velocity in cm/sec with which the fish head moved from its initial to final position during the C-start reaction, calculated as (HDD/C-sT). All parameters were collected from each trial and averaged across the two trials.

**Fig 1.**
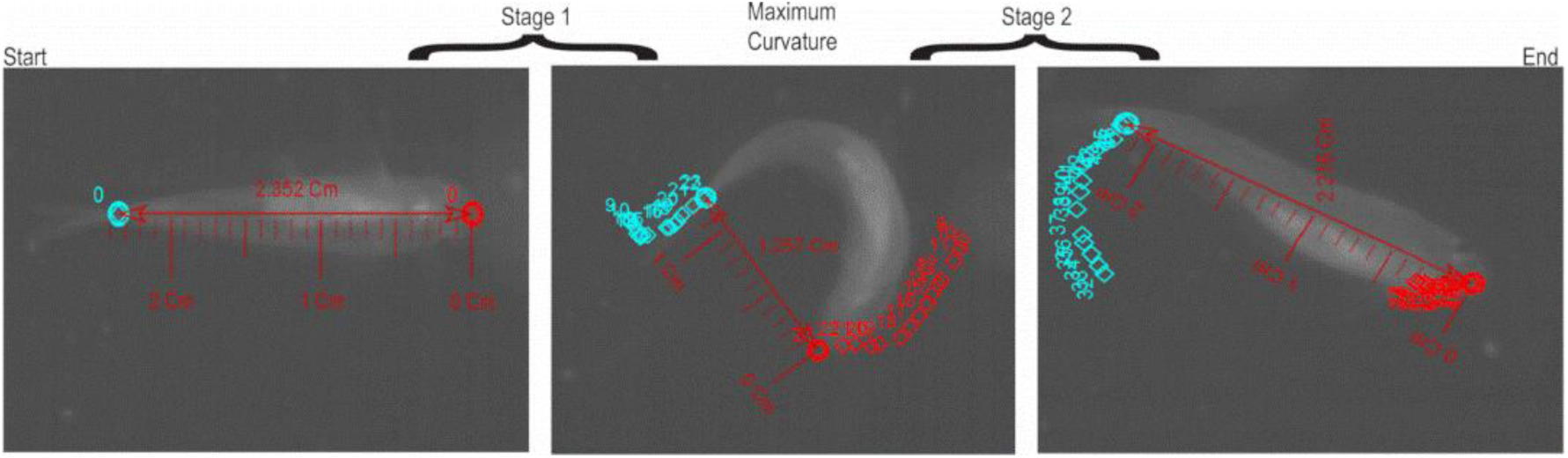
C-start escape response. From left to right: stage one, maximum curvature, stage 2. Red points track head movement, blue points track tail movement. Maximum Curvature is defined by the smallest distance between the head and tail during the C-start response.

Each performance parameter was assessed for outliers, which were defined as values exceeding 1.5 times the interquartile range from the first or third quartiles. Individuals were excluded from the analyses if any one of their performance parameters fell outside this range, ensuring that the dataset reflected typical performance trends without the influence of extreme values. Out of the 280 fish that survived to testing, 22 fish were identified as outliers and removed from the data set. Outliers did not show significant bias toward fish with anomalies, as confirmed by a chi-squared analysis (Table A in S1 Appendix).

### Vertebral Anomaly Identification

The four anomaly classes used in this study included vertebral compressions, fusions, spinal curvatures, and haemal and neural spine malformations (defined by any anomalies (e.g., splitting, bending) of haemal or neural parts of the vertebrae). Identification of compressions and fusions followed Witten et al. (2009) (Fus = types 6-8, Co = types 1-5). Anomalies were recorded from analog x-rays taken at the Field Museum in Chicago, IL. Two individuals assessed anomalies present for each fish to ensure proper identification. Micro-CT scans were taken to produce better visuals for anomaly figures but not used for the identification of the anomalies (Fig 2). Vertebral variables were quantified from x-rays, including total vertebral count and the vertebral ratio (the ratio of precaudal to caudal vertebrae). The Weberian apparatus and urostyle were excluded from counts and anomalies in these structures were not included in this study.

**Fig 2.**
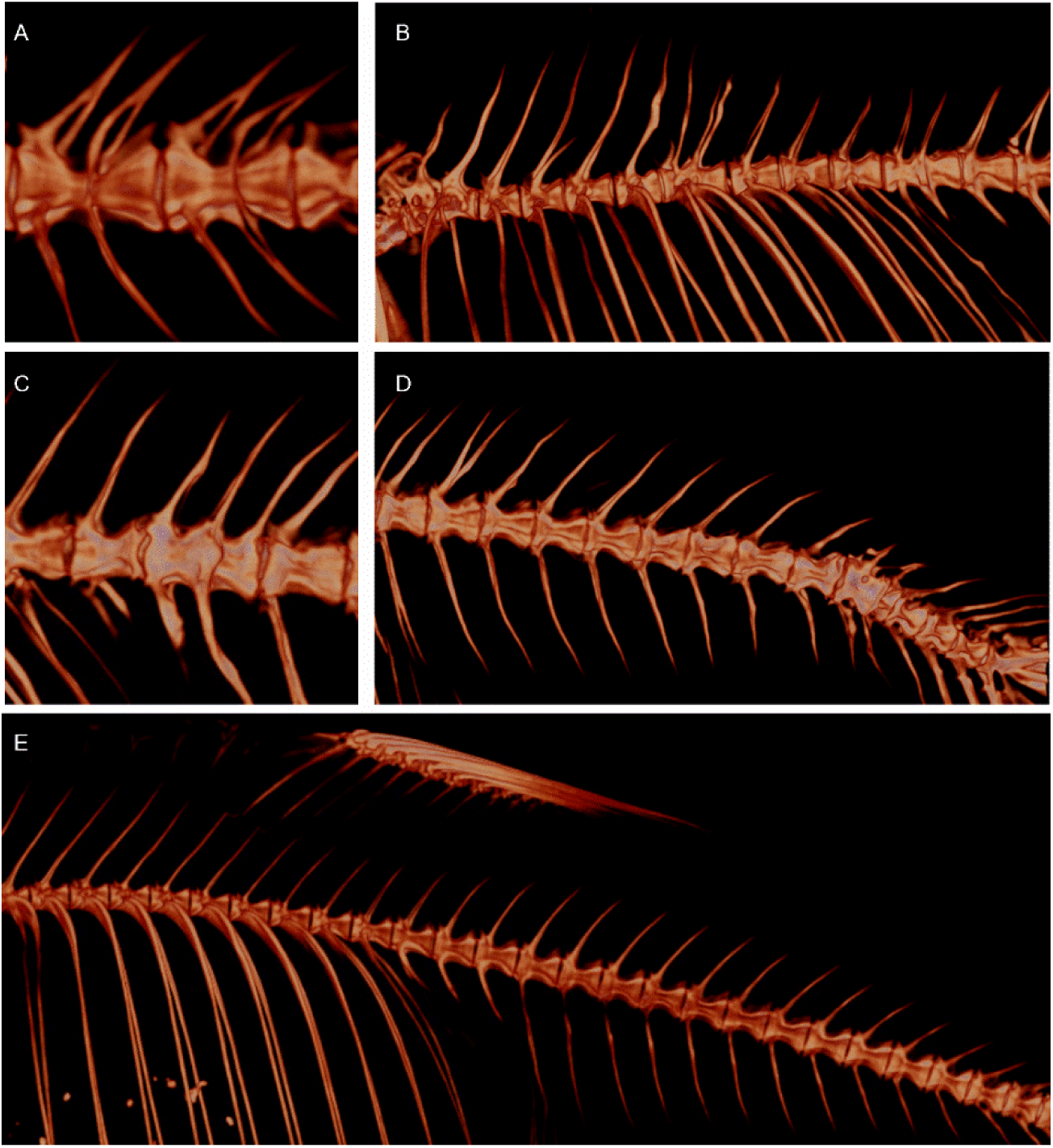
Micro-CT scans of *A. mexicanus* vertebrae with anomalies. A) Fusion. B) Haemal/Neural C) Compression and D) Curvature. E) Normal *A. mexicanus* spine.

Precaudal vertebrae were defined by the presence of ribs, while caudal vertebrae were identified by the presence of both haemal and neural spines.

## Data Analysis

All statistical analyses were using R programming software, versions 4.2.2 and 4.2.3.

### Examining the Influence of Standard length Across Treatments and Anomaly Classes

Previous research has shown that SL can be a significant predictor of performance outcomes including velocity and displacement because larger fish often exhibit superior swimming capabilities due to increased muscle mass and hydrodynamic efficiency (15,25,26). One-way ANOVA tests were used to assess the effects of treatment and the number of anomaly types on mean SL. Additionally, Welch two-sample t-tests were conducted to compare the variation in SL between groups with and without specific anomaly types. These tests assess whether there are significant differences in SL between the groups since this could influence performance.

### Evaluation of the Frequency of Vertebral Anomalies Across Treatments

To evaluate whether the occurrence of vertebral anomalies varied across treatment groups, chi-squared tests were employed. These tests were used to determine if the distribution of anomalies was independent of the treatment conditions. The vertebral anomalies analyzed included the general absence or presence of deformities, as well as specific types such as fusions, compressions, curvatures, haemal and neural spine anomalies, and severity of deformities. For each variable, contingency tables were generated to summarize the frequency of anomalies across treatment groups. Chi-squared tests of independence were applied to these tables to assess whether the observed frequencies differed significantly from the expected frequencies under the assumption that the treatment and the presence of anomalies are independent.

In cases where the assumptions of the chi-squared test were violated, such as when expected variable counts were low, Monte Carlo simulations containing 10,000 iterations were conducted to provide more accurate p-values. This approach provided estimates for the significant relationships between treatment groups and the occurrence of anomalies.

### Assessing Combined Effects on C-Start via MANCOVA

Multivariate analysis of covariance (MANCOVA) was employed to determine the combined effects of multiple independent variables on a suite of dependent variables that quantitatively describe the C-start swimming performance. Swimming parameters were used as the dependent variables for our analysis including average C-start time (C-sT), average displacement velocity (DV), average head displacement distance (HDD), and average curvature coefficient (CC). These variables collectively reflect the diverse aspects of C-start swimming performance influenced by vertebral deformities. The independent variables in the MANCOVA model were treatment, absence/presence of anomalies, haemal/neural anomalies (HN), fusions (Fus), compressions (Co), curvatures (Cu), and type count (the total number of anomaly types). The covariates included SL, precaudal/caudal vertebral ratio, total vertebral count, and total anomaly count. The significance of these effects was evaluated using Pillai’s trace, which is robust to violations of assumptions such as homogeneity of variances and covariances (27).

### GLMM & LMM Analysis of C-Start and Vertebral Anomalies

The goal for these analyses was to assess the interdisciplinary effects of anomalies on C-start performance. A generalize linear mixed model (GLMM) was used for binary outcomes such as the presence or absence of each anomaly type. This approach allows us to handle binary data appropriately by using a logit link function as part of the binomial family of distributions. In this model, ‘Cross’ was included as a random effect to accommodate genetic relatedness while treatment conditions, SL, total count of vertebrae and the ratio of precaudal to caudal vertebrae were incorporated as fixed effects.

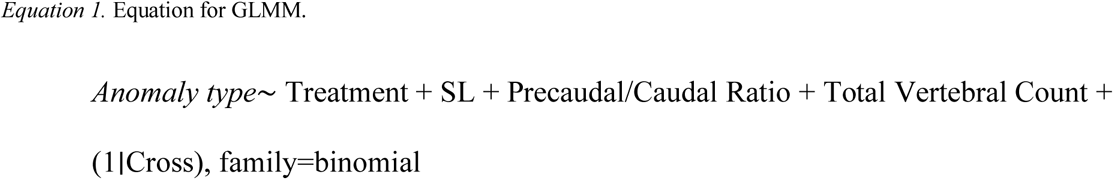

Model validation involved diagnostic checks for normality of residuals and homogeneity of variances. Normality was assessed visually through Q-Q plots, and homogeneity was evaluated using residual vs. fitted value plots. Additionally, we checked for potential overdispersion in the GLMM and model stability, ensuring that no individual level had an excessive influence on the model’s estimates.

Linear Mixed Models (LMMs) were performed to evaluate the effects vertebral anomalies had on C-start escape performances of *A. mexicanus*. The hierarchical structure of our data necessitated the use of mixed-effects models. Specifically, we included the variable ‘Cross’ as a random effect to account for genetic relatedness among individual specimens, thereby capturing the non-independence of data points within the same genetic lineage.

The first model integrated a suite of predictors as fixed effects, including treatment conditions, standard length (SL), total count of vertebrae, and the ratio of precaudal to caudal vertebrae.

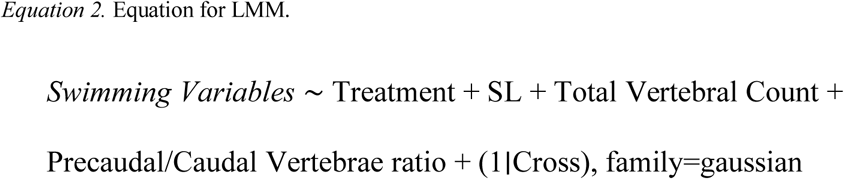

The second LMM focused exclusively on the specific types of vertebral anomalies. This model considered the standard length and deformity types as a fixed effect and ‘Cross’ as a random effect to control for genetic factors.

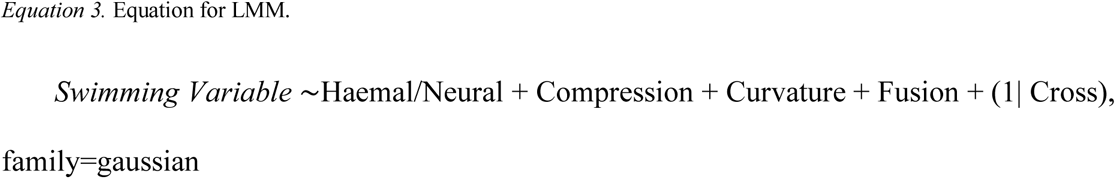

The third and final LMM focused on the total impact anomalies may have on performance, including the total count of anomalies as well as the number of types of anomalies present as fixed effects, again with Cross used as the random effect, see Equation 4.

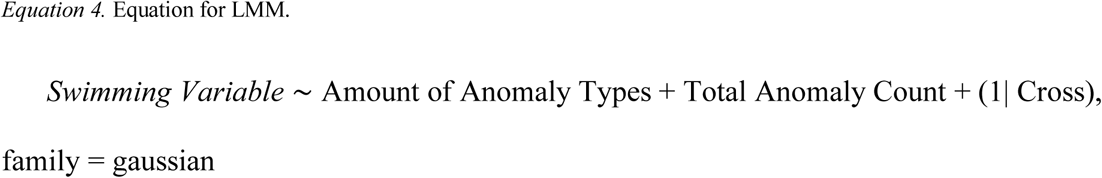

## Results

Out of the 300 expected fish, 258 specimens were used in this study. Early on in the rearing process, all individuals from the short-term temperature treatment in cross 1 died, as well as a total of 5 fish in the long term treatment. In total, cross 1 only had 15 fish, with 10 being in the constant temperature treatment and 5 being in the long-term treatment. In addition to these losses, cross 2 had one less individual in its long-term treatment, resulting in total of 9, cross 12 had one less specimen in the constant and short-term treatment, and cross 21 had one less specimen in the constant and short-term treatment. In addition, 22 fish were removed from this experiment due to one or more swimming parameters falling outside of the 1.5 interquartile range used to identify outliers.

Vertebral anomalies were common among the specimens. Of the 258 specimens used in this study, 161 fish (62.4% of fish in the experiment) had at least one anomaly. Haemal and neural anomalies were the most common anomalies (46.9% of total specimens, 75.2% of specimens with anomalies) and were more frequent than the number of fish who do not have anomalies (37.6%) in each treatment (Fig 3).

**Fig 3.**
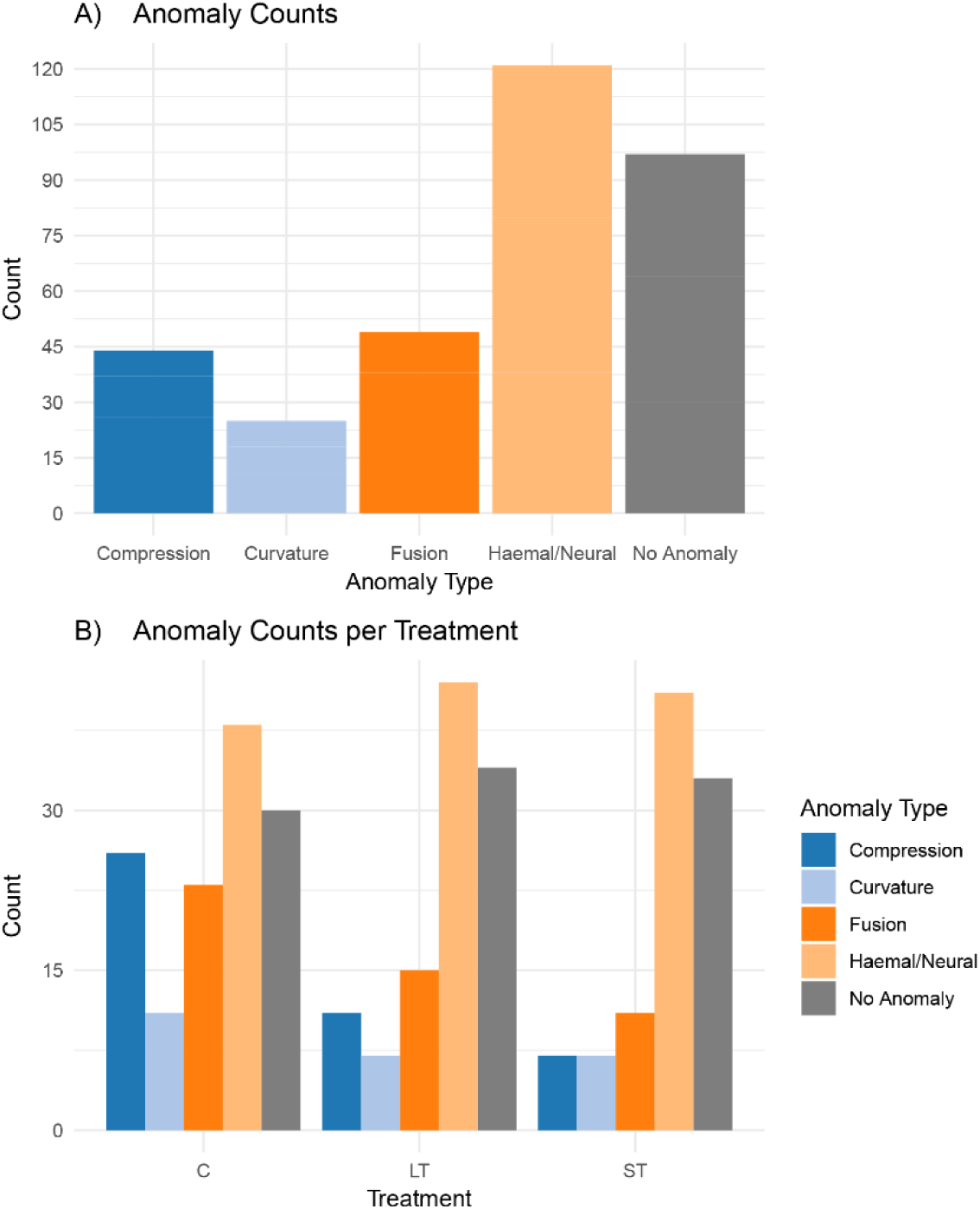
Bar graphs showing distribution of anomalies. A) Total anomalies count overall. B) Total anomalies count per Treatment.

### Standard Length Across Treatments and Anomalies

The results of the one-way ANOVA showed that treatment had significant effects of SL (df = 2, f = 10.39, p = 4.16E-05, Table B in S1 Appendix) while the number of anomaly types did not (df = 4, f = 2.082, p = 0.0836, Table C in S1 Appendix; full Tukey HSD results in Table E in S1 Appendix). A Tukey HSD post-hoc test showed that the LT treatment fish had significantly greater SL than both the ST (p = <0.001) and C (p = 0.012) treatments (Table D in S1 Appendix). There was no significant difference detected between the ST and C treatments (p = 0.226) (Table D in S1 Appendix). The t-test comparing SL between groups with and without anomalies indicated that there was no significant difference between groups (Table 1, Table F in S1 Appendix), with only the general presence/absence of anomalies nearing a significant result (df =196.65, p = 0.09343). Significant differences in SL were also analyzed for the crosses, with an ANOVA showing that cross did have significant effects on SL (df = 9, f = 2.258, p = 0.0191, Table G in S1 Appendix), although the follow up Tukey HSD test showed that the only crosses that differed significantly in SL were between crosses 18 and 1 (p = 0.0431) and crosses 18 and 2 (p = 0.0318) (Fig 4, Table E in S1 Appendix).

**Fig 4.**
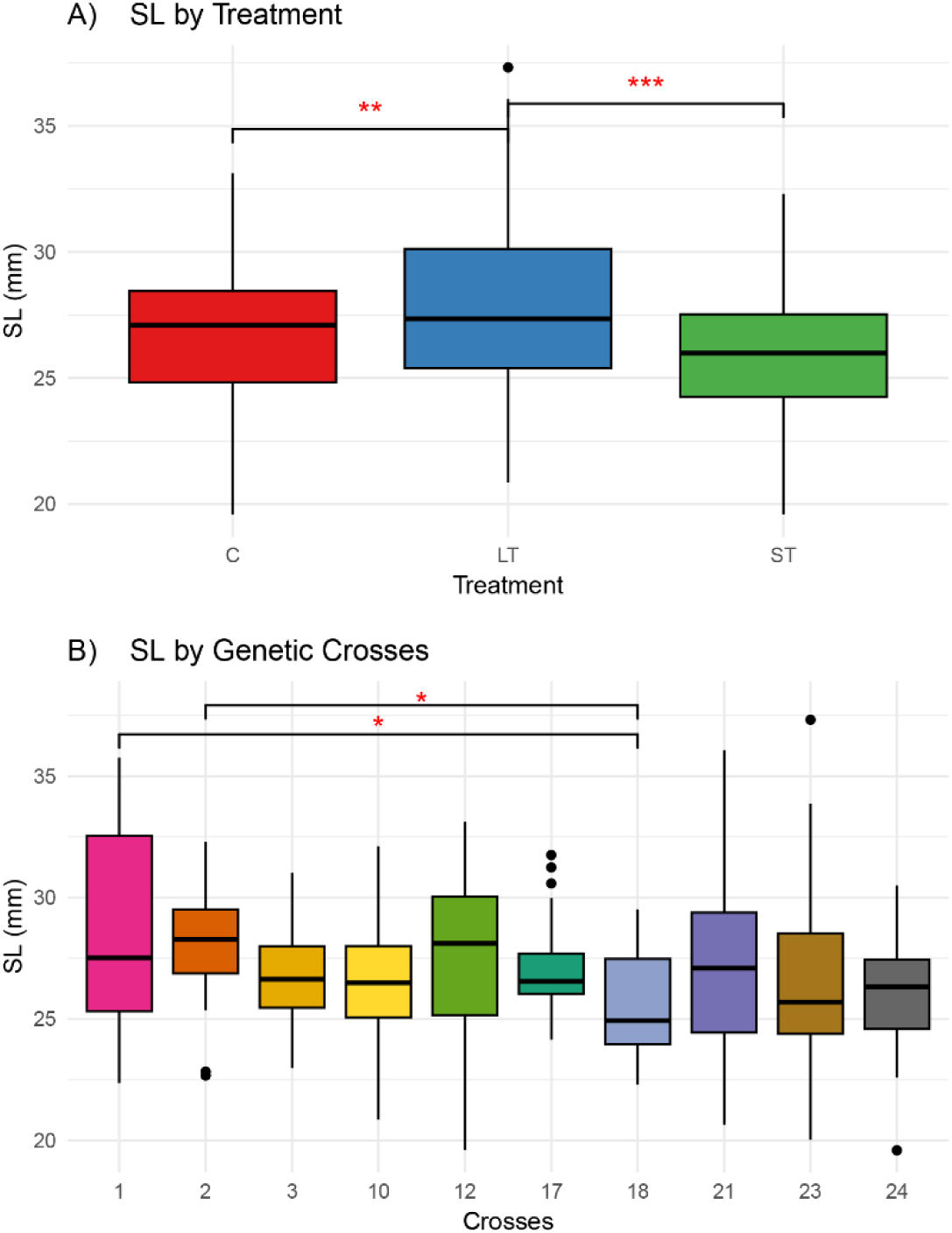
Boxplots showing SL. A) SL across Treatments, and B) SL across Crosses. Asterisks (*) indicate significance (* = 0.05, ** ≤ 0.01, *** ≤ 0.001).

**Fig 5.**
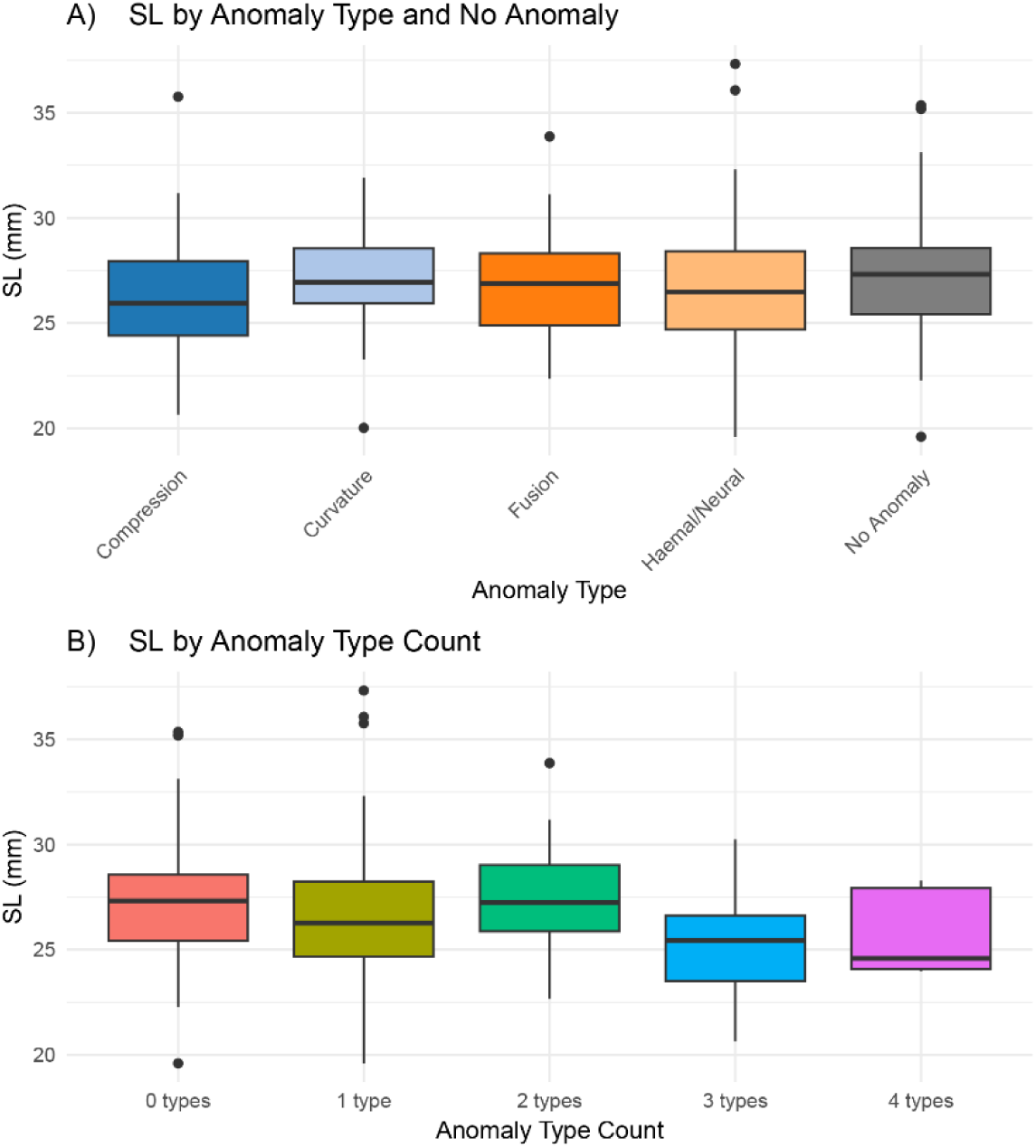
Boxplots showing the range of SL. A) SL for each anomaly types B) SL for the anomaly type count.

**Table 1.** T-test results testing significant differences in SL among anomaly variables.

| T-test results table | t | df | p |
| --- | --- | --- | --- |
| Presence/Absence Anomalies | 1.6857 | 196.65 | 0.09343 |
| Haemal/Neural | 0.67195 | 253.77 | 0.5022 |
| Compression | 1.5857 | 63.174 | 0.1178 |
| Curvature | 0.99689 | 29.574 | 0.3269 |
| Fusion | 1.5937 | 80.623 | 0.1149 |

Full table available as Table E in S1 Appendix.

### The Frequency of Vertebral Anomalies Across Treatments

Monte Carlo chi-squared tests were performed to assess whether there is a significant association between treatment groups and the presence of various vertebral anomalies. This approach was used to test for the distribution of anomaly types across different treatments, allowing for robust p-value estimation even in cases of small sample sizes or expected counts. The results indicate a significant difference in compression anomalies (X² = 15.84, p = <0.0001) and a nearly significant difference in fusion anomalies (X² = 5.2059, p = 0.06849) among treatments. Curvatures (X² = 1.3127, p = 0.5191) and haemal/neural (X² = 0.55175, p = 0.7617) anomalies did not display statistically significant differences among the groups. Likewise, the number of different types of anomalies did not differ significantly among groups (X² = 12.473, p = 0.1247), nor did total count of anomalies (X² = 33.194, p = 0.4064), see Table 2.

**Table 2.** Monte Carlo Chi-squared analysis.

| Monte Carlo Chi-Squared table |  |  |
| --- | --- | --- |
| | $X^2$ | p-value |
| Absence/Presence of Anomalies | 1.0525 | 0.5797 |
| Compression | 15.838 | **<0.0001** |
| Fusion | 5.2059 | 0.06849 |
| Curvature | 1.3127 | 0.5191 |
| Haemal/Neural | 0.55175 | 0.7617 |
| Anomaly Type Count | 12.473 | 0.1247 |
| Total Anomaly Count | 33.194 | 0.4064 |

Assessing combined effects on C-Start via MANCOVA was conducted to examine the effects of environmental and anatomical factors on total C-start escape performance in fish (Table 3). The results indicate that treatments significantly impacted performance metrics (p = 0.002), while fusion anomalies show a marginally non-significant effect (p = 0.069). Other anomaly types did not have significant or near-significant effects on total performance. Among the covariates, SL had the strongest influence (p = 1.43E-09), followed by total vertebral count (p = 0.007) and the precaudal/caudal vertebral ratio (p = 0.018), both of which also significantly affected performance (Table 3).

**Table 3.** MANCOVA table.

| MANCOVA | formula = manova(Performance ~ Treatment + Absence/Presence of Anomalies + HN + Fusion + Compression+ Curvature + Type Count + SL + Precaudal/Caudal Ratio + Total Vertebral Count + Total Anomaly Count) |  |  |  |
| --- | --- | --- | --- | --- |
|  | Df | Pillai | Approx F | Pr(>F) |
| Treatment | 2 | 0.081835 | 3.4415 | **0.002445** |
| Deform | 1 | 0.008799 | 0.7132 | 0.544972 |
| Haemal/Neural | 1 | 0.009429 | 0.7647 | 0.514824 |
| Fusion | 1 | 0.028942 | 2.3943 | 0.069001 |
| Compression | 1 | 0.000901 | 0.0724 | 0.974699 |
| Curvature | 1 | 0.01487 | 1.2126 | 0.305756 |
| Type Count | 3 | 0.040652 | 1.1127 | 0.350996 |
| SL | 1 | 0.166969 | 16.1017 | **1.43E-09** |
| Precaudal/Caudal Ratio | 1 | 0.040936 | 3.4288 | **0.017785** |
| Total Vertebral Count | 1 | 0.048437 | 4.0892 | **0.007417** |
| Total Anomaly Count | 1 | 0.007434 | 0.6016 | 0.614513 |
| Residuals | 244 |  |  |  |

Independent variables include Treatment, the general presence of anomalies, the presence of specific types of anomalies, and the count of how many types are present. Covariates are shown in gray and include the SL, precaudal/caudal vertebral ratio, total vertebral count and total anomaly count. Dependent variables are performance parameters.

### GLMM Analysis of C-Start and Vertebral Anomalies

A generalized linear mixed model (GLMM) was employed to evaluate the significance of various treatment and anatomical parameters in predicting the occurrence of different types of anomalies in fish. The model incorporated fixed effects for treatment conditions, specifically long-term (LT) and short-term (ST) temperature treatments, SL, precaudal/caudal ratio, and total vertebral count. The random effect of ‘Cross’ was included to account for variability among different related groups of fish. Given that the response variable, anomaly presence/absence, is binary (indicating the presence or absence of each type of anomaly), the binomial family was used in the GLMM. Table 4 presents the p-values, outlining the statistical significance of each predictor for the various anomaly types.

**Table 4.** GLMM p-values for predicting the presence of anomaly types based on temperature treatment, standard length, vertebrae ratio, and total vertebral count as fixed effects and cross as the random effect.

| GLMM p-values |  | glmer(Anomaly Presence/Absence ~Treatment + SL +<br>Precaudal/Caudal Ratio+ Total Vertebral Count + (1\|Cross) ,<br>family = binomial) |  |  |  |  |
| --- | --- | --- | --- | --- | --- | --- |
|  | Intercept | Treatment -<br>LT | Treatment –<br>ST | Vertebral<br>Count | Precaudal/Caudal<br>Ratio | Standard<br>Length |
| Haemal/Neural | **0.0449** | 0.1566 | 0.1913 | **0.0138** | 0.0963 | 0.6578 |
| Compression | 0.18017 | **0.03406** | **0.00292** | 0.49875 | 0.15261 | 0.10094 |
| Curvature | 0.791 | 0.386 | 0.296 | 0.783 | 0.929 | 0.323 |
| Fusion | 0.256 | 0.449 | 0.112 | 0.421 | 0.41 | 0.167 |

Full table available in supplemental materials, Table I in S1 Appendix.

GLMM analysis revealed that fish raised under LT treatment (est. = -0.893, p = 0.03406) and ST treatment (est. = -1.469, p = 0.00292) developed significantly fewer compression anomalies compared to those in treatment C (Table 4). This indicated that elevated temperatures during any amount of time in early development did not contribute to the presence of compressions, with significantly more compressions observed in fish reared in treatment C (Fig 3b). Additionally, the occurrence of haemal/neural anomalies was significantly influenced by the total vertebral count (est. = -0.4319, p = 0.0138), indicating that fish with a lower number of vertebrae were more likely to develop these anomalies. Interestingly, a significant result with the occurrence of haemal/neural anomalies was indicated in the intercept (est. = 10.78, p = 0.0449), indicating that this specific anomaly type may develop not only due to low vertebral count, but potentially another variable not measured in this study. The precaudal/caudal vertebral ratio also showed a marginally non-significant effect on haemal/neural anomalies (est. = 3.674, p = 0.0963), suggesting that this ratio may be associated with the development of haemal/neural anomalies, though the relationship is not statistically conclusive (Table 4).

### LMM Analysis of C-Start and Vertebral Anomalies

We applied LMM to assess the influence of several predictors on swimming performance parameters and the presence of vertebral anomalies in *A. mexicanus*. Two LMM were performed to address the main questions of this study, including if C-start performance parameters were influenced by anomaly type, number of different anomalies present, and/or total anomaly counts (Tables 5 & 6). Another LMM was also done to assess other potential influences on swimming performance variation, including temperature treatments, total vertebral count, vertebrae ratio, and standard length, Table 7.

The first LMM analysis looked at the effects anomaly types had on the swimming performance parameters (Table A in S1 Appendix1; summarized in Table 5). Out of all four parameters measured, only CC was shown to have any significant interactions with anomalies. Specifically, haemal/neural and fusion anomalies were found to significantly impact curving ability. Haemal/neural anomalies had a negative effect on the CC (est. = -0.00229, t = -2.44, p = 0.0153), indicating a decreased curving ability. Conversely, fusion anomalies had a positive effect on CC (est. = 0.00293, t = 2.351, p = 0.0195), demonstrating an increased curving ability. Thus, CC is disproportionately impacted by anomalies more than any of the other C-start performance parameters, reflected in Fig 6.

**Fig 6.**
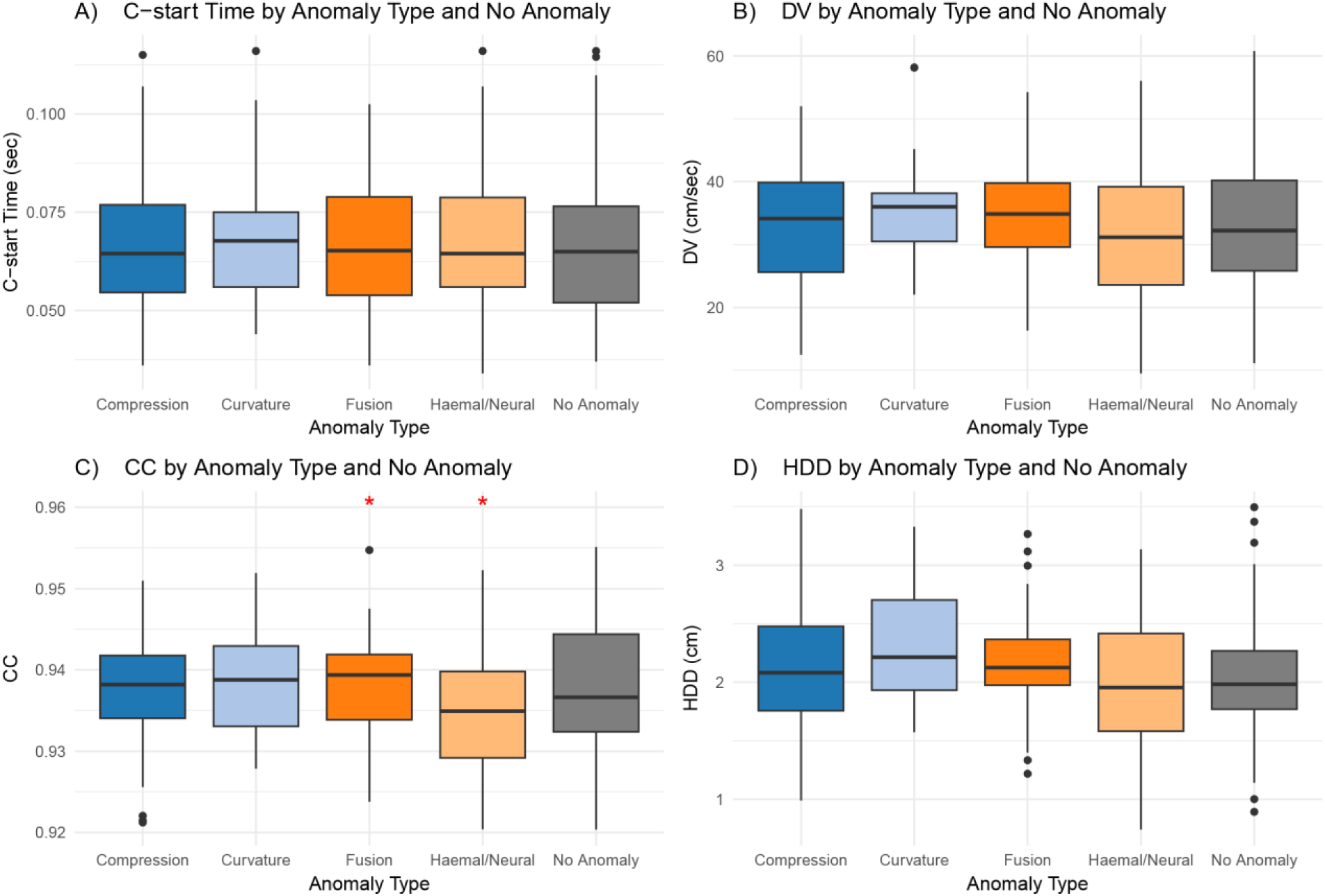
Boxplot showing how each anomaly type and absence of anomalies impact swimming performance parameters. A) C-start Time B) DV C) CC D) HDD. Red asterisks (*) indicate significance (* = 0.05, ** ≤ 0.01, *** ≤ 0.001).

**Table 5.** Linear Mixed Model p-values in terms of anomaly types and standard length as fixed effects with cross as a random effect.

| LMM p-values |  | lmer(formula = Swimming parameters ~ Haemal/Neural + Compression + Curvature + Fusion + (1\| Cross), family = gaussian) |  |  |  |
| --- | --- | --- | --- | --- | --- |
|  | Intercept | Haemal/Neural | Compression | Curvature | Fusion |
| C-Start Time | <2e-16 | 0.429 | 0.927 | 0.59 | 0.546 |
| DV | 5.87E-14 | 0.13 | 0.945 | 0.249 | 0.245 |
| HDD | <2e-16 | 0.571 | 0.945 | 0.238 | 0.157 |
| CC | <2e-16 | **0.015** | 0.7968 | 0.1084 | **0.0195** |

Full table available in supplemental materials, Table K in S1 Appendix.

The second LMM considered both the number of anomaly types and the total count of vertebrae with anomalies for each individual (Table 6). For C-start time, the amount of anomaly types and the total number of vertebral anomalies had no significant effect indicating that the time of the C-start response was not significantly influenced by the presence of vertebral anomalies. However, a positive trend was in the boxplots, where fish with all 4 anomaly types were shown to have comparatively lower and therefore faster C-start time than all other categories (Fig 7). For HDD, neither the number of anomaly types nor the total anomaly count had a significant impact. However, the presence of all four anomaly types had a significant positive effect on DV (est. = 10.6818, t = 2.206, p = 0.0283) and CC (est. = 0.007551, t = 2.00, p = 0.0466), resulting in faster velocity speeds and increased curving ability, although the total anomaly count did not have a significant impact (Fig 7). Similarly, fish with the four anomaly types experienced a positive impact in curving ability during the c-start response, with the total anomaly count also having a positive but marginally non-significant effect on CC (est. = 0.007551, t = -1.662, p = 0.0978) (Fig 7).

**Fig 7.**
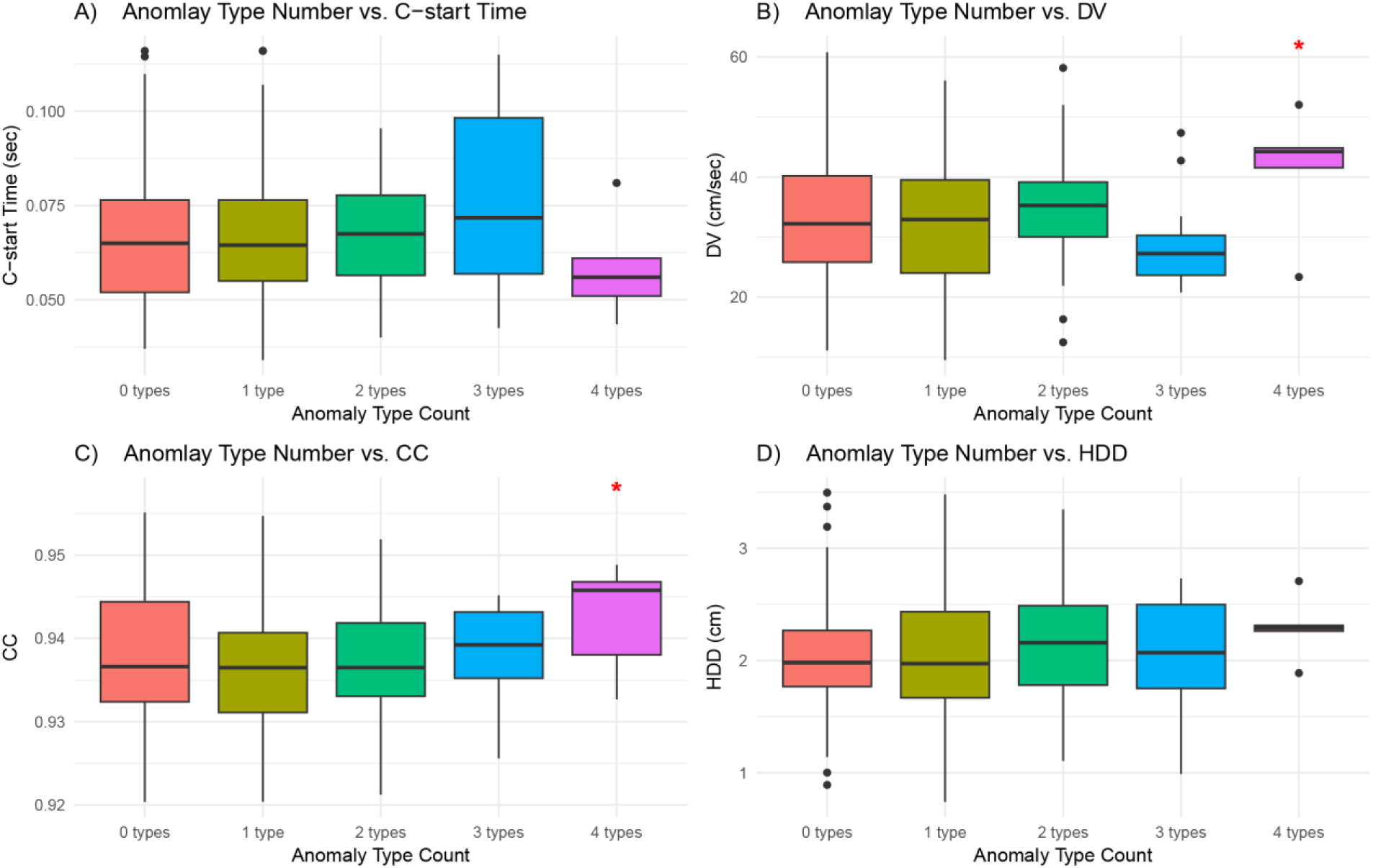
Boxplots showing the results from Table 7. Anomaly Type Counts indicate the number of specific anomaly types present, 0 types indicate no anomaly types are present, while 4 types indicate all anomaly types are present. A) C-start Time, B) DV, C) CC, D) HDD. Red asterisks (*) indicate significance (* = 0.05, ** ≤ 0.01, *** ≤ 0.001).

**Table 6.**
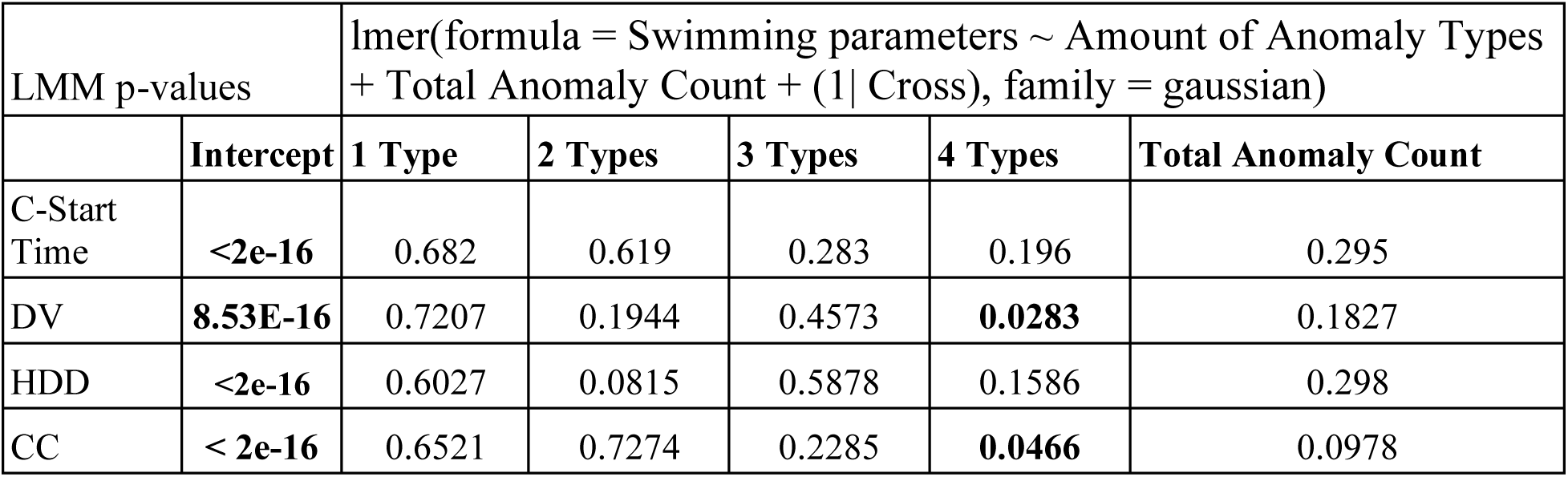
Linear Mixed Model p-values for performance parameters in terms of number of anomaly types present and total anomaly count, with cross as a random effect.

**Table 7.**
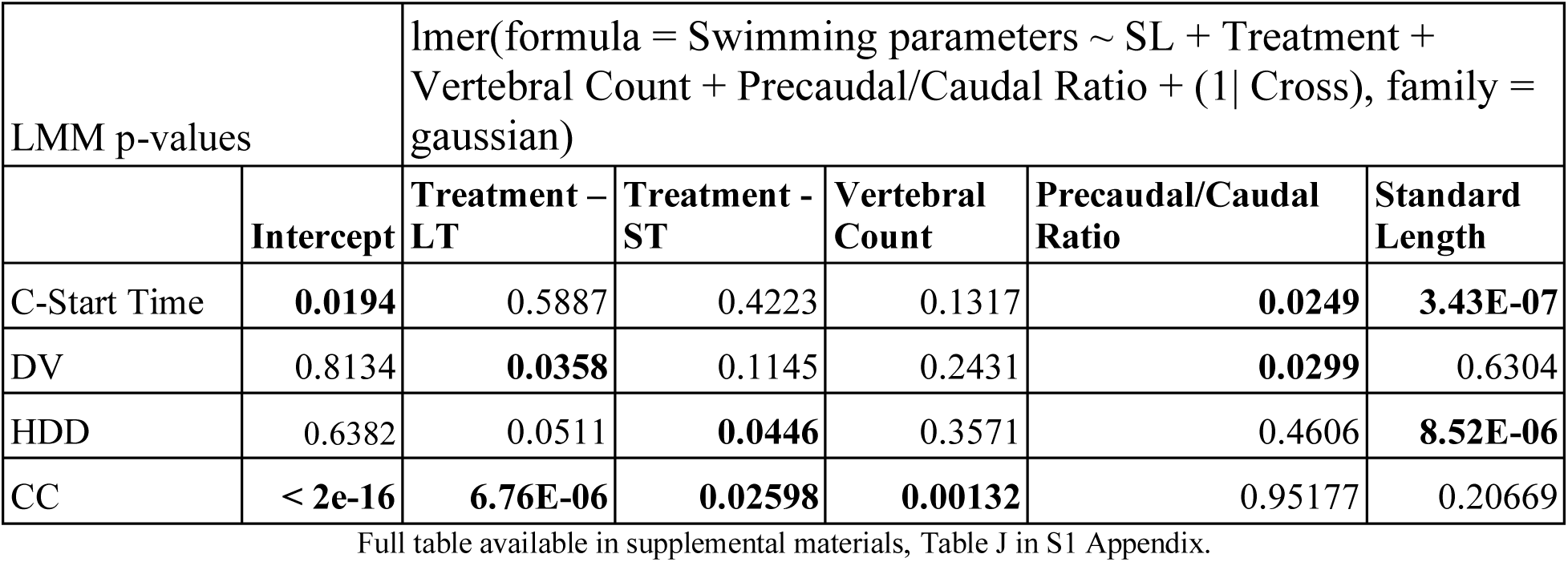
Linear Mixed Models p-values for performance parameters in terms of standard length, treatments, total vertebral count, and vertebral ratio as fixed effects and cross as a random effect.

| LMM p-values |  | lmer(formula = Swimming parameters ~ SL + Treatment + Vertebral Count + Precaudal/Caudal Ratio + (1\| Cross), family = gaussian) |  |  |  |  |
| --- | --- | --- | --- | --- | --- | --- |
|  | Intercept | Treatment – LT | Treatment - ST | Vertebral Count | Precaudal/Caudal Ratio | Standard Length |
| C-Start Time | **0.0194** | 0.5887 | 0.4223 | 0.1317 | **0.0249** | **3.43E-07** |
| DV | 0.8134 | **0.0358** | 0.1145 | 0.2431 | **0.0299** | 0.6304 |
| HDD | 0.6382 | 0.0511 | **0.0446** | 0.3571 | 0.4606 | **8.52E-06** |
| CC | **< 2e-16** | **6.76E-06** | **0.02598** | **0.00132** | 0.95177 | 0.20669 |

Full table available in supplemental materials, Table J in S1 Appendix.

The influence of SL and the Precaudal/Caudal ratio on performance metrics showed mixed effects (Table 7). SL had a positive effect on C-start time (est. = 0.002014, p < 0.001), meaning that longer fish took more time to complete a response. Additionally, SL was positively associated with HDD (est. = 0.05087, p < 0.001), but it had no significant effect on DV or CC (see Figure A in S1 Appendix). Vertebral counts in our specimens ranged from 27 to 32, with most individuals exhibiting an average of 30 vertebrae. Although a few individuals fell slightly outside the typical species range of 29–31 vertebrae, the majority were within this expected range. A higher precaudal/caudal ratio had a negative effect on C-start time (est. = -0.03991, p = 0.0249), indicating that fish with a more equal number of precaudal and caudal vertebrae exhibited faster escape response times. A higher precaudal/caudal ratio also had a positive effect on DV (est. = 22.7116, p = 0.0299), meaning fish with a closer ratio of vertebrae type moved quicker during their escape. Vertebral count significantly impacted CC (est. = 0.001981, t = .331, p = 0.00132), showing that fish with more vertebrae were able to achieve higher curvatures during the response (Fig 8).

**Fig 8.**
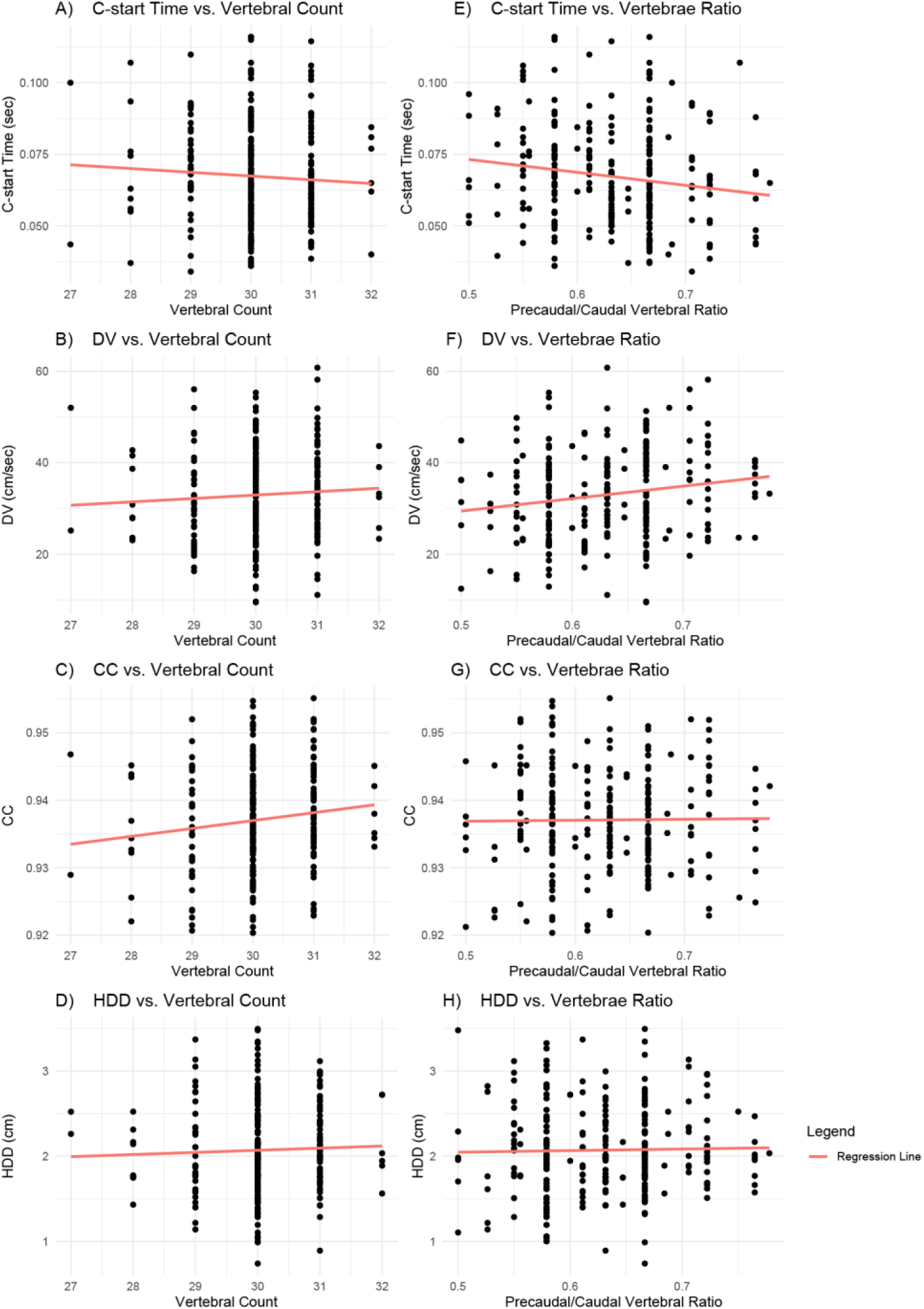
Scatter plot shows the relationship between performance metrics vertebral count, and precaudal/caudal ratio. Linear regression line shown in red. A-D shows vertebral count, E-H shows precaudal/caudal ratio. A/E) C-start Time, B/F) DV, C/G) CC, D/H) HDD.

Regarding treatment effects, the LT treatment significantly reduced DV (est. = -3.2483, t = -2.11, p = 0.0358) and CC (est. = -0.00529, t = -4.598, p < 0.001), suggesting slower speeds and reduced curving ability (Table 7). The ST treatment had significant negative effects on HDD (est. = -0.1669, t = -2.019, p = 0.0446) and on CC (est. = -0.002653, t = -2.24, p = 0.026), indicating fish in ST treatment covered less distance and had less curving ability (Table 7). Overall CC was negatively impacted by both temperature treatments, indicating that long and short periods of elevated temperatures during development can negatively impact a fish’s ability to bend its body during the C-start reaction (Fig 9).

**Fig 9.**
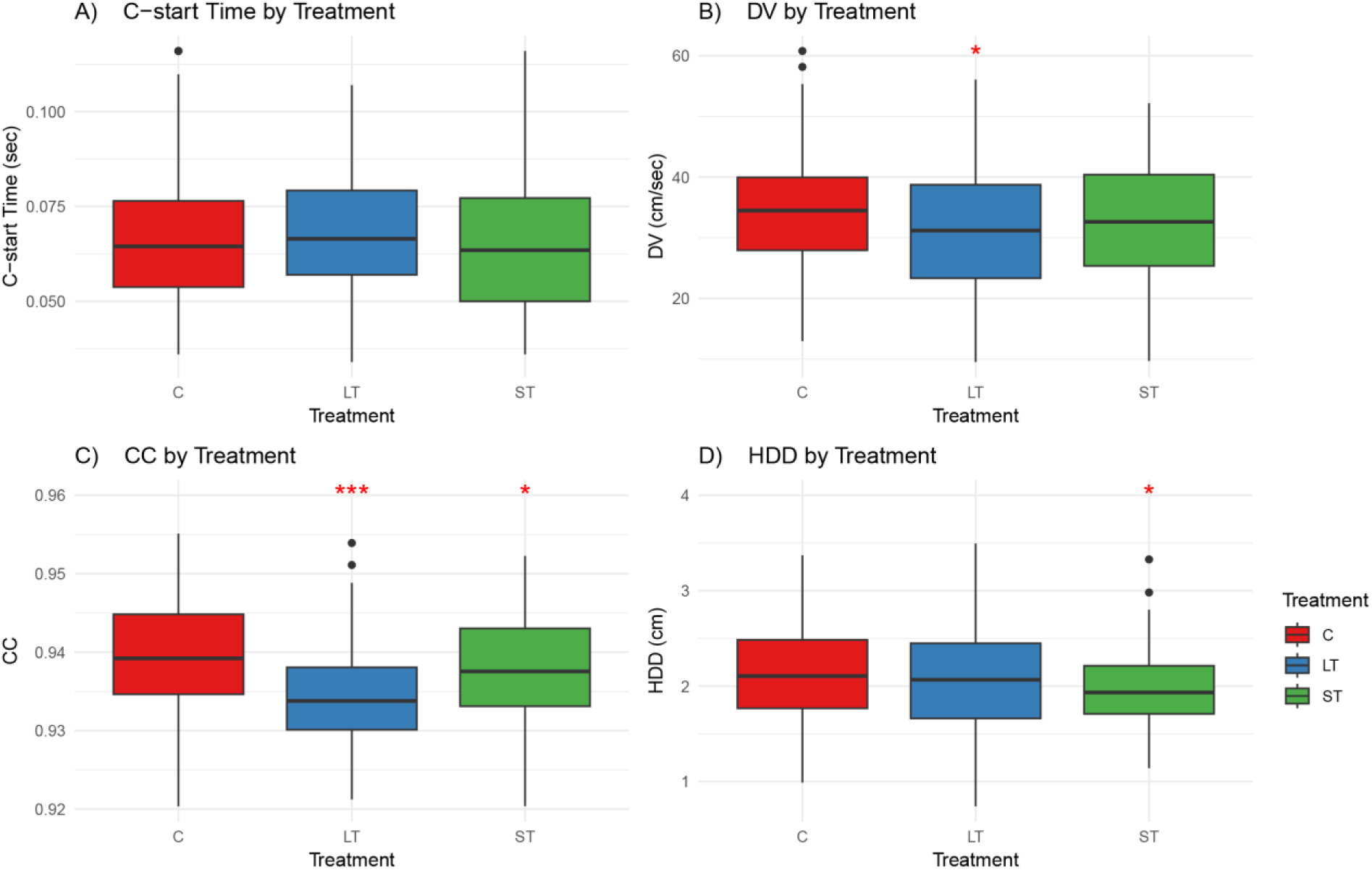
Boxplot showing C-start performance metrics across treatments. A) C-start Time B) DV, C) CC, D) HDD. Red asterisk(s) indicate significant difference. Black dots show spread of data for each treatment (* = 0.05, ** ≤ 0.01, *** ≤ 0.001).

Across all LMM, a few common trends emerged. Treatment conditions, particularly the LT treatment, consistently reduced DV and CC, indicating slower speeds and less curvature during the escape response (Table 7). The ST treatment had a negative effect on CC and HDD, showing that fish reared with a short period of elevated temperature during early development had a reduced curving ability and covered less distance while escaping. CC and DV were positively affected when all four anomaly types were present, while time and distance did not significantly change. SL had a positive effect on both c-start time and HDD, meaning that although longer fish took more time to escape, they were able to travel farther distances. Precaudal/caudal ratio showed positive effects on performance where a closer ratio of precaudal/caudal vertebrae improved both C-start time and DV, indicating that the reactions were performed faster. CC was the most affected performance metric, being sensitive to temperature treatments, total vertebral count, and the presence of all four anomaly types, and being directly impacted by fusion and haemal/neural anomalies.

## Discussion

Vertebral anomalies in *Astyanax mexicanus* had complex and varied impacts on swimming performance metrics. While some anomalies significantly affected specific aspects of C-start performance, their effects were generally weaker than those of overall body size and vertebral anatomy. Haemal/neural spine anomalies had a negative effect on the CC, and increasing anomaly severity (the total number of anomalies) reduced DV and CC (Table 6, Fig B in S1 Appendix). In contrast, vertebral fusions and the presence of all four anomaly types were associated with improved performance metrics (Table 6, Fig 7). The swimming performance metrics most closely associated with fitness, C-start time and HDD (14), were unaffected by anomaly type or severity but were both significantly influenced by SL and vertebral morphology (Table 7, Fig 8). Experimental temperature treatments also impacted C-start performance metrics in addition to the development of certain anomalies, which highlights the role environmental conditions play in shaping locomotion (28–30). Overall, these results demonstrate that the relationship between vertebral anomalies, morphology, and performance is highly context dependent.

### The Impact of Vertebral Anomalies on C-start Performance

While skeletal anomalies are generally thought to impair locomotive abilities (6,12,31,32), our study revealed a complex interaction between vertebral deformities and swimming performance. Notably, DV and CC were both significantly positively impacted when all four anomaly types were present (Table 6). However, a non-significant negative trend was found when considering the total number of vertebral anomalies (severity) (Table 6, Figure B in S1 Appendix). These results suggest that a greater total number of anomalies negatively affects speed (DV) and curving ability (CC), but having all four anomaly types does not necessarily produce the same negative effect (Fig 7). Simply put, the variety of anomaly types was associated with improved performance, whereas the overall number of anomalies present showed a negative trend. A possible explanation is that the presence of multiple anomaly types does not necessarily indicate more severe skeletal deformities. Fish may exhibit substantial resilience to skeletal anomalies when it comes to their burst-swimming performance. This is consistent with an increasing number of reports of wild-caught sexually mature fish that exhibit skeletal anomalies, challenging assumptions about the detrimental effects of such deformities on survival mechanisms like escaping predators (1,7,10,11,43,44). Other studies have shown that the relationship between skeletal anomalies and swimming performance can be complex. Research on *Rhamdia quelen* showed that pollutant exposure during early development can increase the prevalence of skeletal anomalies without completely impairing escape performance (33). Another study looking at hatchery reared lumpfishes showed that although vertebral anomalies impacted the dorsal and lateral body shape of the fish, the deformities did not significantly impact swimming performance alone and that body shape changes may have indirectly impacted swimming performance (34).

When investigating the specific impacts anomalies had on the performance parameters, we found the CC was the only parameter significantly affected by individual anomaly types, although fusions had a marginally non-significant negative impact on HDD during the escape response. Interestingly, fusions seemed to have a significant positive effect on CC while haemal/neural anomalies had a significant negative impact (Table 5) indicating that fish with fusions had a greater curving ability while fish with haemal/neural anomalies had a reduced curving ability, impacting the “C” formation critical for the escape (13,15,35). Fusions have been found to have variable and indirect impacts on swimming performance parameters, with current research suggesting that fusions may impact body shape, maneuverability, and energy expenditure, which could be related to adaptive muscle compositions (34,36–40). Studies in cleaner fish have also revealed that skeletal anomalies, including fusions, can occasionally yield unexpected performance trade-offs (41). The positive association between vertebral fusions and curving ability suggests that the biomechanical consequence of anomalies may be more complex than traditionally assumed and may involve compensatory changes in muscle function had on curving ability, consistent with the role of flexibility in enhancing locomotor diversity (35).

These findings align with studies showing that mechanical stress and functional modularity can influence biomechanical performance during development, as observed in fish adapting to various environmental conditions (42). Similar results have been found in Powell et al. (2009), where scoliotic fish were found to have reduced performance in reaching critical swimming speeds, while fish with compressions performed on par with individuals who did not present any anomalies (12).

### The Effects of Other Anatomical Traits on C-start Performance

SL significantly influenced C-start time and HDD, with larger fish exhibiting more effective escape responses (Table 7, Fig 8), consistent with the well-established relationship between body size and swimming performance (12,45,46). Larger individuals may compensate for potential mechanical disadvantages associated with vertebral anomalies through greater muscle mass and energy reserves (14,25,46,47). Vertebral count also significantly influenced performance, specifically CC, highlighting the functional importance of vertebral anatomy in locomotion (36,44,48). Overall, general anatomical traits including SL, precaudal/caudal vertebral ratio, and vertebral count explained more variation in C-start escape performance than vertebral anomalies themselves (Table 3, Fig. 8). These traits directly influence force transmission, body flexibility, and swimming mechanics emphasizing the importance of anatomy and body condition in determining escape performance (13,36,49,50).

### Temperature Effects on C-start Performance and Anomaly Development

Environmental conditions not only influence anatomical development but also escape response performance (16,52). Compression and fusion anomalies were particularly sensitive to developmental temperature (Table A in S1 Appendix). Chi-squared analyses indicated significant differences among treatments, and GLMM results further demonstrated significant association between temperature treatment and vertebral compressions (Table 4). Fish reared in elevated temperatures displayed significantly fewer compression anomalies. This is consistent with findings in *Sparus aurata* larvae, where early temperature exposure significantly shaped skeletal development and swimming performance (16), suggesting that changes in developmental temperature significantly influence the occurrence of anomalies (3,39,52,53).

Along with the chi-squared analysis, a LMM (Table 5, Fig 9) also demonstrated significant effects temperature treatments had on C-start performance, with LT treatment significantly reducing DV, causing slower speeds, and both LT and ST treatments having negative impacts on CC. This shows that fish in these treatments had reduced curving ability and speed (Fig 9), which would negatively impact two major components of C-start responses (14,15). The variability in anomaly occurrence and performance outcomes reflects broader trends in phenotypic disparity driven by environmental factors during developmental windows (54). These results demonstrate that changes in developmental temperature can influence both the occurrence of vertebral anomalies, like compressions and fusions, and key swimming performance measures, such as DV and CC. Overall, our results show that temperature-sensitive anomaly development aligns with evidence that developmental plasticity can have lasting effects on performance, particularly under changing environmental conditions. Previous research has also shown that some of these effects could be epigenetically heritable, causing a transgenerational impact on future fish populations in the wild (52).

### Broader Implications

These results are particularly relevant for Neotropical conservation efforts, given the region’s high biodiversity and the environmental pressures it faces. Neotropical freshwater ecosystems are home to an extraordinary variety of fish species, many of which rely on effective C-start escape responses to evade predators (18,55,56). Although vertebral anomalies affected specific components of escape performance, particularly DV and CC, their effects on C-start time and HDD were limited. The fitness consequences of vertebral anomalies may be more subtle than previously thought and may depend on specific anomaly type, severity, and environmental conditions experienced during development (1,12,57). This finding is especially relevant as human-induced environmental changes, such as pollution and climate change, increasingly alter developmental conditions for aquatic species (7,10,51).

Furthermore, environmental stressors like pollution and habitat loss exacerbate skeletal anomalies in Neotropical fish, threatening the regions aquatic biodiversity and fish fitness, consistent with broader patterns of anthropogenic impacts (1,2,4,58,59). The sensitivity of compression and fusion anomalies to temperature raises concerns about how climate change could exacerbate skeletal deformities in some fish species. As global temperatures change, the developmental conditions that lead to skeletal deformities may become more prevalent (16,36,60,61). This could have cascading effects on fish survival, since even small effects can reduce their ability to evade predators and thrive.

Reducing anthropogenic stressors, maintaining optimal growth conditions, and protecting natural habitats will be essential to ensure the long-term survival of Neotropical fish populations (55,56,62). Preserving the developmental synchrony and mitigating anthropogenic stressors could reduce the prevalence of skeletal anomalies (6,63). By addressing these environmental stressors and their links to skeletal and vertebral anomalies, conservation efforts can focus on reducing the factors that contribute to these deformities. This approach can be key to preserving the unique biodiversity of Neotropical ecosystems as they continue to face mounting pressures from anthropogenic factors.

This study underscores the importance of re-evaluating the ecological ramifications of vertebral anomalies within the context of fish population dynamics, performance, and conservation strategies. While anomalies are typically viewed under a negative lens due to their perceived impact on fish marketability and welfare in aquaculture settings (6,8,61,64,64,65), their influence in natural populations might be less straightforward. More research is needed to understand their role in the resilience and adaptability of fish species facing environmental pressures.

## Acknowledgements

We thank the staff of the Research Support Facility at DePaul University for their assistance with the care of *Astyanax mexicanus* throughout this project. We also thank The Field Museum for access to its X-ray imaging resources, which were essential for identifying and documenting vertebral anomalies. We are grateful to Dr. Kory Evans at Rice University for providing access to the micro-CT scanner used to generate 3D scans of vertebral anomalies for the figures. Finally, we thank DePaul University for supporting this research from 2021 to 2023.

## Supporting Information

**S1 Appendix.** Supplementary tables and figures. Supplementary statistical analyses and figures examining vertebral anomalies, developmental conditions, morphology, and C-start escape performance in Astyanax mexicanus.

## Notes

### Competing Interest Statement

The authors have declared no competing interest.

### Summary of Updates

Updated to remove format issue from document.

