## Supplementary Materials for "The impact of temperature-induced vertebral anomalies on C-start swimming performance in Astyanax mexicanus (Teleostei: Characidae)"

S.1 Table

Chi-squared analysis to check if there is a significant difference with and without the outliers. Chi-squared table.

| Chi-squared Test for Association Between Vertebral Anomalies and Outlier Status in Fish | |
|---|---|
| X-squared | 0.53824 |
| df | 1 |
| p-value | 0.4632 |

S.2 Table

ANOVA Results for SL by Treatment

| ANOVA Results for SL by Treatment | | | | | |
|---|---|---|---|---|---|
| **Factor** | **Df** | **Sum Sq** | **Mean Sq** | **F value** | **p-value** |
| Treatment | 2 | 163.1 | 81.56 | 10.39 | 4.61e-05* |
| Residuals | 255 | 2002.3 | 7.85 | | |

S.3 Table

ANOVA Results for SL by Anomaly Type Count

| ANOVA Results for SL by Anomaly Type Count | | | | | |
|---|---|---|---|---|---|
| **Factor** | **Df** | **Sum Sq** | **Mean Sq** | **F value** | **p-value** |
| AnomalyTypeNum | 4 | 69 | 17.251 | 2.082 | 0.0836 |

| | | | |
|---|---|---|---|
| Residuals | 253 | 2096 | 8.286 |

S.4 Table

Tukey HSD Pairwise Comparisons for SL by Treatment

| Tukey HSD Pairwise Comparisons for SL by Treatment | | | | |
|---|---|---|---|---|
| **Comparison** | **Difference** | **Lower CI** | **Upper CI** | **p-value** |
| LT - C | 1.2198 | 0.2182 | 2.2214 | 0.0123 |
| ST - C | -0.708 | -1.7185 | 0.3025 | 0.226 |
| ST - LT | -1.9278 | -2.9383 | -0.9172 | 0.00003* |

S.5 Table

Tukey HSD Pairwise Comparisons for SL by Anomaly Type Count

| Tukey HSD Pairwise Comparisons for SL by Anomaly Type Count | | | | |
|---|---|---|---|---|
| **Comparison** | **Difference** | **Lower CI** | **Upper CI** | **p-value** |
| 1 type - 0 types | -0.6207 | -1.7296 | 0.4881 | 0.5387 |
| 2 types - 0 types | 0.0611 | -1.4984 | 1.6206 | 0.9999 |
| 3 types - 0 types | -1.9772 | -4.2384 | 0.284 | 0.118 |
| 4 types - 0 types | -1.5009 | -5.1279 | 2.1262 | 0.7868 |
| 2 types - 1 type | 0.6819 | -0.8582 | 2.2219 | 0.7418 |

| | | | | |
|---|---|---|---|---|
| 3 types - 1 type | -1.3564 | -3.6042 | 0.8914 | 0.4619 |
| 4 types - 1 type | -0.8801 | -4.4989 | 2.7386 | 0.963 |
| 3 types - 2 types | -2.0383 | -4.5393 | 0.4628 | 0.1688 |
| 4 types - 2 types | -1.562 | -5.3432 | 2.2192 | 0.7878 |
| 4 types - 3 types | 0.4763 | -3.6442 | 4.5968 | 0.9978 |

S.6 Table

T-Test Results for Standard Length (SL) by Anomaly Type

| **Anomaly Type** | **t-value** | **df** | **p-value** | **Mean (Group 0)** | **Mean (Group 1)** | **95% Confidence Interval (Lower)** | **95% Confidence Interval (Upper)** |
|---|---|---|---|---|---|---|---|
| General Presence of Deformity | 1.6857 | 196.65 | 0.09343 | 27.28196 | 26.64963 | -0.10742 | 1.37208 |
| Compression | 1.5857 | 63.174 | 0.1178 | 27.01449 | 26.26909 | -0.19389 | 1.68468 |
| Fusion | 1.5937 | 80.623 | 0.1149 | 27.01493 | 26.34327 | -0.16694 | 1.51027 |
| Curvature | 0.99689 | 29.574 | 0.3269 | 26.94562 | 26.3444 | -0.63121 | 1.83366 |
| Haemal/Neural | 0.67195 | 253.77 | 0.5022 | 27.00139 | 26.75826 | -0.46942 | 0.95566 |

S.7 Table

ANOVA Results for Standard Length (SL) by Cross

| Factor | Df | Sum Sq | Mean Sq | F value | p-value |
|---|---|---|---|---|---|
| Cross | 9 | 164 | 18.23 | 2.258 | 0.0191* |

| | | | |
|---|---|---|---|
| Residuals | 248 | 2001 | 8.07 |

S.8 Table

Tukey HSD Pairwise Comparisons for SL by Cross

| Comparison | Difference | Lower CI | Upper CI | p-value |
|---|---|---|---|---|
| Significant Pairwise Comparisons ($p < 0.05$) | | | | |
| 18 - 1 | -3.295 | -6.539 | -0.051 | 0.043 |
| 18 - 2 | -2.585 | -5.054 | -0.117 | 0.032 |
| Non-Significant Comparisons ($p > 0.05$) | | | | |
| 2 - 1 | -0.71 | -3.954 | 2.534 | 0.999 |
| 3 - 1 | -2.024 | -5.252 | 1.203 | 0.598 |
| 10 - 1 | -2.152 | -5.397 | 1.092 | 0.517 |
| 12 - 1 | -1.163 | -4.488 | 2.162 | 0.983 |
| 17 - 1 | -1.655 | -4.883 | 1.572 | 0.828 |
| 21 - 1 | -1.484 | -4.696 | 1.727 | 0.901 |
| 23 - 1 | -1.764 | -4.961 | 1.433 | 0.758 |
| 24 - 1 | -2.61 | -5.837 | 0.618 | 0.232 |
| 3 - 2 | -1.314 | -3.761 | 1.132 | 0.786 |
| 10 - 2 | -1.442 | -3.911 | 1.026 | 0.692 |
| 12 - 2 | -0.453 | -3.027 | 2.121 | 0.999 |
| 17 - 2 | -0.945 | -3.392 | 1.501 | 0.966 |
| 21 - 2 | -0.774 | -3.2 | 1.651 | 0.991 |
| 23 - 2 | -1.054 | -3.46 | 1.352 | 0.927 |

| | | | | |
|---|---|---|---|---|
| 24 - 2 | -1.9 | -4.346 | 0.547 | 0.285 |
| 10 - 3 | -0.128 | -2.574 | 2.318 | 1 |
| 12 - 3 | 0.861 | -1.691 | 3.414 | 0.986 |
| 17 - 3 | 0.369 | -2.055 | 2.793 | 1 |
| 18 - 3 | -1.271 | -3.717 | 1.176 | 0.817 |
| 21 - 3 | 0.54 | -1.863 | 2.943 | 0.999 |
| 23 - 3 | 0.26 | -2.123 | 2.643 | 1 |
| 24 - 3 | -0.585 | -3.009 | 1.839 | 0.999 |
| 12 - 10 | 0.989 | -1.584 | 3.563 | 0.967 |
| 17 - 10 | 0.497 | -1.95 | 2.943 | 0.999 |
| 18 - 10 | -1.143 | -3.612 | 1.326 | 0.9 |
| 21 - 10 | 0.668 | -1.758 | 3.093 | 0.997 |
| 23 - 10 | 0.388 | -2.018 | 2.794 | 1 |
| 24 - 10 | -0.457 | -2.904 | 1.989 | 1 |
| 17 - 12 | -0.492 | -3.045 | 2.06 | 1 |
| 18 - 12 | -2.132 | -4.706 | 0.442 | 0.203 |
| 21 - 12 | -0.321 | -2.854 | 2.211 | 1 |
| 23 - 12 | -0.601 | -3.115 | 1.912 | 0.999 |
| 24 - 12 | -1.447 | -3.999 | 1.106 | 0.729 |
| 18 - 17 | -1.64 | -4.086 | 0.807 | 0.501 |
| 21 - 17 | 0.171 | -2.232 | 2.574 | 1 |
| 23 - 17 | -0.109 | -2.492 | 2.274 | 1 |
| 24 - 17 | -0.954 | -3.378 | 1.47 | 0.962 |

| | | | | |
|---|---|---|---|---|
| 21 - 18 | 1.811 | -0.615 | 4.236 | 0.34 |
| 23 - 18 | 1.531 | -0.875 | 3.937 | 0.578 |
| 24 - 18 | 0.686 | -1.761 | 3.132 | 0.997 |
| 23 - 21 | -0.28 | -2.642 | 2.082 | 1 |
| 24 - 21 | -1.125 | -3.528 | 1.278 | 0.893 |
| 24 - 23 | -0.845 | -3.229 | 1.538 | 0.981 |

S.9 Table

GLMM Full Table

| **Model** | **Effect Type** | **Effect** | **Estimate/Variance** | **Std. Error/Std. Dev.** | **z-value** | **Pr(>\|z\|)** |
|---|---|---|---|---|---|---|
| **Haemal/Neural** | Fixed Effects | **(Intercept)** | **10.78369** | **5.37785** | **2.005** | **0.0449** |
| | | TreatmentLT | 0.47512 | 0.33542 | 1.417 | 0.1566 |
| | | TreatmentST | 0.44956 | 0.34407 | 1.307 | 0.1913 |
| | | SL | -0.02042 | 0.04611 | -0.443 | 0.6578 |
| | | PreC_Caud_Ra | 3.67368 | 2.20889 | 1.663 | 0.0963 |
| | | **TotalVertCount** | **-0.4319** | **0.17531** | **-2.464** | **0.0138** |
| | Random Effect | Cross (Intercept) | 0.0188 | 0.1371 | | |
| **Compression** | Fixed Effects | (Intercept) | 8.884 | 6.62864 | 1.34 | 0.18017 |
| | | **TreatmentLT** | **-0.89329** | **0.42148** | **-2.119** | **0.03406** |
| | | **TreatmentST** | **-1.46942** | **0.49372** | **-2.976** | **0.00292** |
| | | SL | -0.10641 | 0.06487 | -1.64 | 0.10094 |
| | | PreC_Caud_Ra | -4.28794 | 2.99778 | -1.43 | 0.15261 |
| | | TotalVertCount | -0.14337 | 0.21194 | -0.676 | 0.49875 |

| | | | | | | |
|---|---|---|---|---|---|---|
| | Random Effect | Cross (Intercept) | 0 | 0 | | |
| **Curvature** | Fixed Effects | (Intercept) | -2.20686 | 8.34666 | -0.264 | 0.791 |
| | | TreatmentLT | -0.46586 | 0.53742 | -0.867 | 0.386 |
| | | TreatmentST | -0.58224 | 0.55665 | -1.046 | 0.296 |
| | | SL | -0.08004 | 0.08101 | -0.988 | 0.323 |
| | | PreC_Caud_Ra | 0.32079 | 3.58165 | 0.09 | 0.929 |
| | | TotalVertCount | 0.0741 | 0.26924 | 0.275 | 0.783 |
| | Random Effect | Cross (Intercept) | 0 | 0 | | |
| **Fusion** | Fixed Effects | (Intercept) | 8.4619 | 7.44615 | 1.136 | 0.256 |
| | | TreatmentLT | -0.32293 | 0.4267 | -0.757 | 0.449 |
| | | TreatmentST | -0.73685 | 0.46364 | -1.589 | 0.112 |
| | | SL | -0.08909 | 0.06442 | -1.383 | 0.167 |
| | | PreC_Caud_Ra | -2.64627 | 3.21156 | -0.824 | 0.41 |
| | | TotalVertCount | -0.19165 | 0.23794 | -0.805 | 0.421 |
| | Random Effect | Cross (Intercept) | 0.5892 | 0.7676 | | |

S.10 Table

LMM of Swimming Performance Parameters with Treatments, SL, Vertebral Count, and Vertebral Ratio

| **Model** | **Effect Type** | **Effect** | **Estimate/Variance** | **Std. Error/Std. Dev.** | **z-value** | **Pr(>\|z\|)** |
|---|---|---|---|---|---|---|

| | | | | | | |
|---|---|---|---|---|---|---|
| **Haemal/Neural** | Fixed Effects | **(Intercept)** | **10.78369** | **5.37785** | **2.005** | **0.0449** |
| | | TreatmentLT | 0.47512 | 0.33542 | 1.417 | 0.1566 |
| | | TreatmentST | 0.44956 | 0.34407 | 1.307 | 0.1913 |
| | | SL | -0.02042 | 0.04611 | -0.443 | 0.6578 |
| | | PreC_Caud_Ra | 3.67368 | 2.20889 | 1.663 | 0.0963 |
| | | **TotalVertCount** | **-0.4319** | **0.17531** | **-2.464** | **0.0138** |
| | Random Effect | Cross (Intercept) | 0.0188 | 0.1371 | | |
| **Compression** | Fixed Effects | (Intercept) | 8.884 | 6.62864 | 1.34 | 0.18017 |
| | | **TreatmentLT** | **-0.89329** | **0.42148** | **-2.119** | **0.03406** |
| | | **TreatmentST** | **-1.46942** | **0.49372** | **-2.976** | **0.00292** |
| | | SL | -0.10641 | 0.06487 | -1.64 | 0.10094 |
| | | PreC_Caud_Ra | -4.28794 | 2.99778 | -1.43 | 0.15261 |
| | | TotalVertCount | -0.14337 | 0.21194 | -0.676 | 0.49875 |
| | Random Effect | Cross (Intercept) | 0 | 0 | | |
| **Curvature** | Fixed Effects | (Intercept) | -2.20686 | 8.34666 | -0.264 | 0.791 |
| | | TreatmentLT | -0.46586 | 0.53742 | -0.867 | 0.386 |
| | | TreatmentST | -0.58224 | 0.55665 | -1.046 | 0.296 |
| | | SL | -0.08004 | 0.08101 | -0.988 | 0.323 |
| | | PreC_Caud_Ra | 0.32079 | 3.58165 | 0.09 | 0.929 |
| | | TotalVertCount | 0.0741 | 0.26924 | 0.275 | 0.783 |
| | Random Effect | Cross (Intercept) | 0 | 0 | | |
| **Fusion** | Fixed Effects | (Intercept) | 8.4619 | 7.44615 | 1.136 | 0.256 |

| | | | | | | |
|---|---|---|---|---|---|---|
| | | TreatmentLT | -0.32293 | 0.4267 | -0.757 | 0.449 |
| | | TreatmentST | -0.73685 | 0.46364 | -1.589 | 0.112 |
| | | SL | -0.08909 | 0.06442 | -1.383 | 0.167 |
| | | PreC_Caud_Ra | -2.64627 | 3.21156 | -0.824 | 0.41 |
| | | TotalVertCount | -0.19165 | 0.23794 | -0.805 | 0.421 |
| | Random Effect | Cross (Intercept) | 0.5892 | 0.7676 | | |

S.11 Table

LMM of Swimming Performance Parameters with Anomaly types.

| Model | Effect Type | Effect | Estimate/Variance | Std. Error/Std. Dev. | df | t-value | Pr(>\|t\|) |
|---|---|---|---|---|---|---|---|
| **C-start Time** | **Fixed Effects** | **(Intercept)** | **6.66E-02** | **1.93E-03** | **18.23** | **34.585** | **<2e-16** |
| | | Haemal/Neural | 0.001832 | 0.002187 | 253 | 0.793 | 0.429 |
| | | Compression | 0.0008684 | 0.002943 | 248.8 | -0.092 | 0.927 |
| | | Curvature | -0.001135 | 0.003632 | 250 | -0.54 | 0.59 |
| | | Fusion | 0.002919 | 0.002897 | 247.4 | 0.605 | 0.546 |
| | Random Effect | Cross (Intercept) | 1.08E-05 | 0.003291 | | | |
| | | Residual | 3.15E-04 | 0.01776 | | | |
| **DV** | Fixed Effects | **(Intercept)** | **33.066** | **1.234** | **14.81** | **26.784** | **5.87E-14** |
| | | Haemal/Neural | -1.867 | 1.23 | 251.88 | -1.518 | 0.13 |
| | | Compression | -0.114 | 1.64 | 247.91 | -0.069 | 0.945 |
| | | Curvature | 2.2342 | 2.029 | 248.21 | 1.155 | 0.249 |
| | | Fusion | 1.907 | 1.6138 | 253 | 1.164 | 0.245 |
| | Random Effect | Cross (Intercept) | 7.826 | 2.798 | | | |

| | | | | | | | |
|---|---|---|---|---|---|---|---|
| | | Residual | 88.328 | 9.398 | | | |
| **HDD** | **Fixed Effects** | **(Intercept)** | **2.05114** | **0.049275** | **26.25** | **41.63** | **<2e-16** |
| | | Haemal/Neural | -0.037749 | 0.066485 | 251.49 | -0.568 | 0.571 |
| | | Compression | 0.006142 | 0.089612 | 250.15 | 0.069 | 0.945 |
| | | Curvature | 0.131002 | 0.110654 | 251.86 | 1.184 | 0.238 |
| | | Fusion | 0.124345 | 0.087477 | 229.45 | 1.421 | 0.157 |
| | Random Effect | Cross (Intercept) | 0.002351 | 0.04849 | | | |
| | | Residual | 0.266828 | 0.51655 | | | |
| **CC** | Fixed Effects | **(Intercept)** | **9.37E-01** | **8.27E-04** | **14.77** | **1133.297** | **<2e-16** |
| | | **Haemal/Neural** | **-2.29E-03** | **9.39E-04** | **252.8** | **-2.44** | **0.0153** |
| | | Compression | -3.24E-04 | 1.26E-03 | 247.8 | -0.258 | 0.7968 |
| | | Curvature | 2.50E-03 | 1.55E-03 | 248.8 | 1.611 | 0.1084 |
| | | **Fusion** | **2.93E-03** | **1.25E-03** | **250.4** | **2.351** | **0.0195** |
| | Random Effect | Cross (Intercept) | 2.50E-06 | 0.001581 | | | |
| | | Residual | 5.20E-05 | 0.007214 | | | |

S.12 Table

LMM of Swimming Performance Parameters with Anomaly Type Count and Total Deformity Count.

| **Model** | **Effect Type** | **Effect** | **Estimate/Variance** | **Std. Error/Std. Dev.** | **df** | **t-value** | **Pr(>\|t\|)** |
|---|---|---|---|---|---|---|---|
| **C-start Time** | **Fixed Effect** | **(Intercept)** | **0.06695** | **0.002103** | **24.4** | **31.832** | **<2e-16** |
| | | 1 type | -0.00119 | 0.002902 | 244.5 | -0.41 | 0.682 |
| | | 2 types | -0.002039 | 0.0041 | 252 | -0.497 | 0.619 |
| | | 3 types | 0.006592 | 0.006122 | 251.4 | 1.077 | 0.283 |
| | | 4 types | -0.01184 | 0.009125 | 249.6 | -1.297 | 0.196 |
| | | Deform_Count | 0.0004099 | 0.0003906 | 252 | 1.049 | 0.295 |
| | Random Effect | Cross (Intercept) | 0.00001173 | 0.003425 | | | |

| | | | | | | | |
|---|---|---|---|---|---|---|---|
| | | Residual | 0.0003102 | 0.017613 | | | |
| **DV** | **Fixed Effect** | **(Intercept)** | **32.91** | **1.3032** | **18.5** | **25.254** | **8.53E-16** |
| | | 1 type | 0.5499 | 1.5361 | 244.14 | 0.358 | 0.7207 |
| | | 2 types | 2.8414 | 2.1838 | 250.47 | 1.301 | 0.1944 |
| | | 3 types | -2.4232 | 3.2547 | 249.04 | -0.745 | 0.4573 |
| | | **4 types** | **10.6818** | **4.8431** | **247.42** | **2.206** | **0.0283** |
| | | Deform_Count | -0.278 | 0.2081 | 250.53 | -1.336 | 0.1827 |
| | **Random Effect** | **Cross (Intercept)** | **7.815** | **2.796** | | | |
| | | Residual | 86.745 | 9.314 | | | |
| **HDD** | **Fixed Effect** | **(Intercept)** | **2.04757** | **0.05456** | **39.73** | **37.53** | **<2e-16** |
| | | 1 type | 0.04435 | 0.08508 | 246.52 | 0.521 | 0.6027 |
| | | 2 types | 0.20826 | 0.11906 | 248.11 | 1.749 | 0.0815 |
| | | 3 types | 0.09674 | 0.17826 | 251.37 | 0.543 | 0.5878 |
| | | 4 types | 0.37665 | 0.26637 | 251.86 | 1.414 | 0.1586 |
| | | Deform_Count | -0.01183 | 0.01134 | 248.04 | -1.043 | 0.298 |
| | Random Effect | Cross (Intercept) | 0.002048 | 0.04525 | | | |
| | | Residual | 0.267514 | 0.51722 | | | |
| **CC** | **Fixed Effect** | **(Intercept)** | **0.9376** | **0.000899** | **20.61** | **1042.89** | **<2e-16** |
| | | 1 type | -0.0005415 | 0.0012 | 243.5 | -0.451 | 0.6521 |
| | | 2 types | 0.0005926 | 0.001698 | 251.9 | 0.349 | 0.7274 |
| | | 3 types | 0.003059 | 0.002534 | 250.7 | 1.207 | 0.2285 |
| | | **4 types** | **0.007551** | **0.003776** | **248.7** | **2** | **0.0466** |
| | | Deform_Count | -0.0002689 | 0.0001618 | 251.9 | -1.662 | 0.0978 |
| | Random Effect | Cross (Intercept) | 0.000002516 | 0.001586 | | | |
| | | Residual | 0.000053 | 0.00728 | | | |

S.13 Figure

Scatter Plots showing the relationship between SL and Swimming Performance Parameters, using the presence/absence of anomalies as a factor. A) C-start Time B) DV C) CC D) HDD.

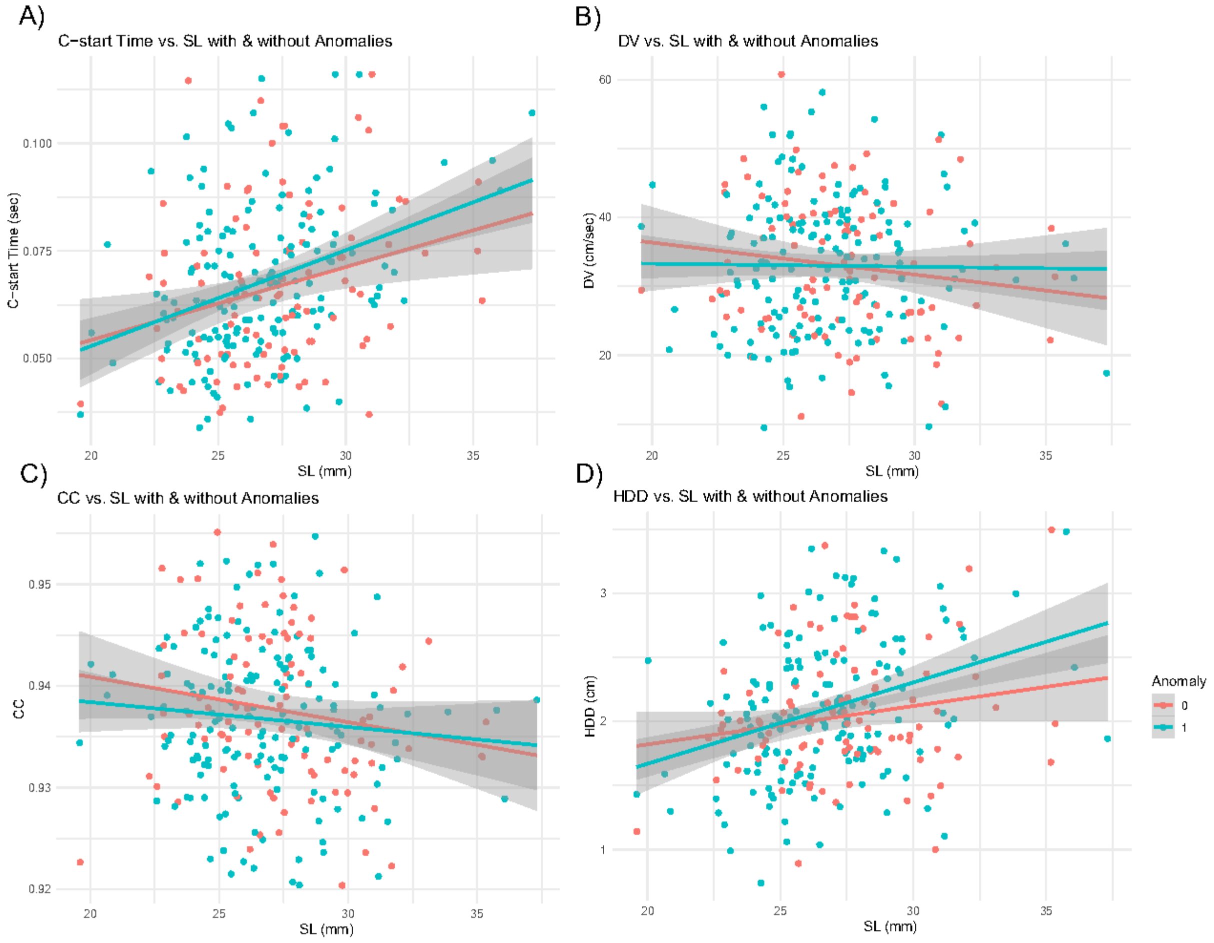

S.14 Figure

Scatter plots showing the relationship between deformity count (severity) and Swimming Performance Parameters. Linear regression line shown in red. A) C-start time, B) DV, C) CC, D) HDD.

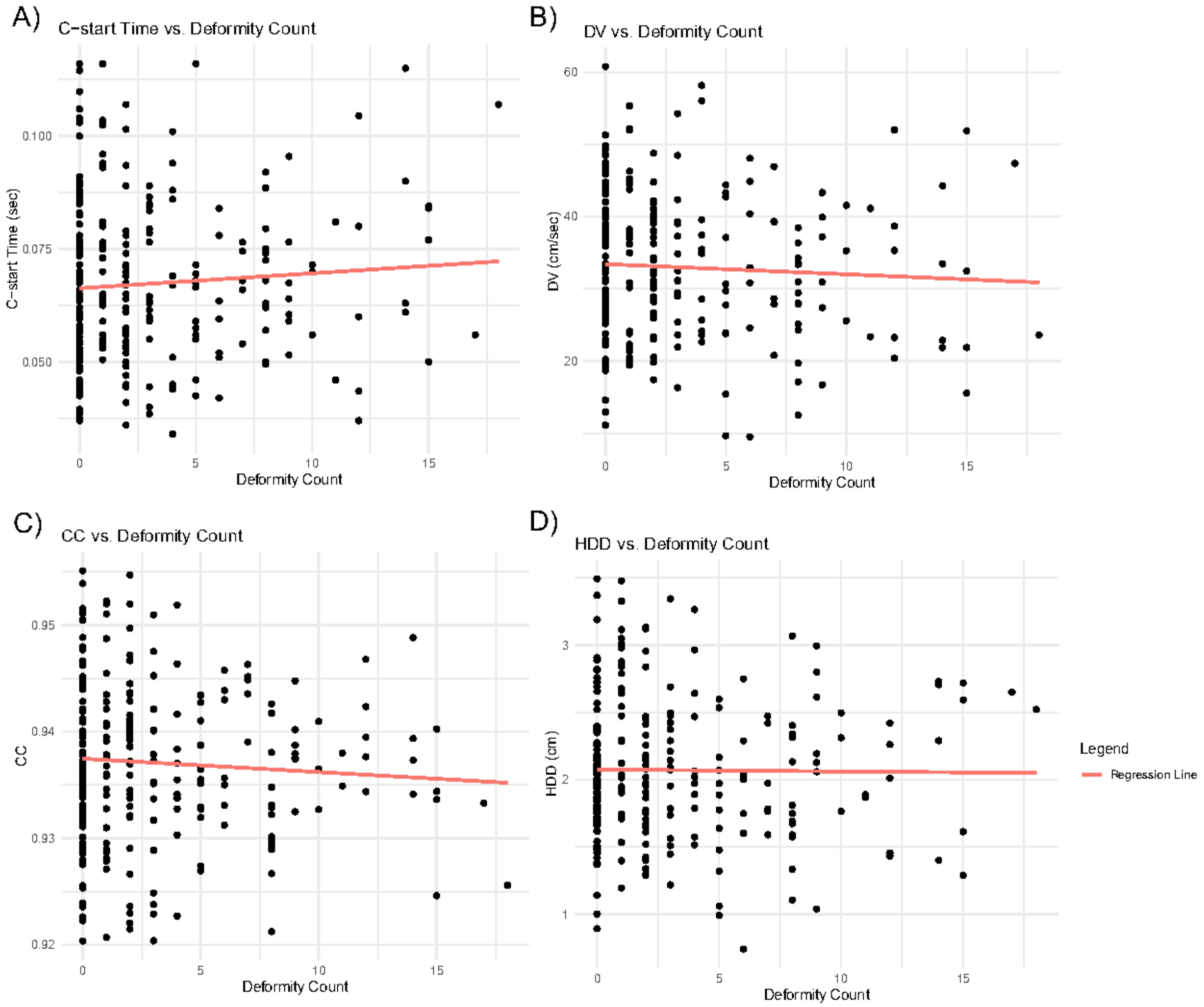
